# Wnt/β-catenin regulates alloreactive T cells for the treatment of hematological malignancies

**DOI:** 10.1101/2021.04.12.439538

**Authors:** Mahinbanu Mammadli, Rebecca Harris, Sara Mahmudlu, Anjali Verma, Adriana May, Rohan Dhawan, Adam T. Waickman, Jyoti Misra Sen, Mobin Karimi

**Affiliations:** Department of Microbiology and Immunology, SUNY Upstate Medical University, Syracuse, NY 13210; National Institute on Aging-National Institutes of Health, 08C218, Biomedical Research Center, 251 Bayview Boulevard, Suite 100, Baltimore, MD 21224; Immunology Program, Department of Medicine, Johns Hopkins School of Medicine, Baltimore, MD 21224

**Keywords:** Key Points: Wnt/β, Wnt/β atenin affects T cell migration to GVHD target organs. atenin affects T cell cytokine production in disease model, Wnt/β-catenin affects T cell gene expression

## Abstract

Allogeneic hematopoietic stem cell transplantation (allo-HSCT) is one of the most widely applied forms of adaptive immunotherapy. Both the detrimental graft-versus-host disease (GVHD) and the beneficial graft-versus-leukemia (GVL) effects occurring after allo-HSCT are largely mediated by alloantigen-reactive donor T cells in the graft. Separating GVHD from GVL effects is a formidable challenge, and a greater understanding of donor T cell biology is required to accomplish the uncoupling of GVHD from GVL. Here, we tested a novel mouse model of β-catenin (*Cat-Tg*) in an allo-HSCT model. Our data show that T cells from *Cat-Tg* mice did not cause GVHD. Surprisingly, *Cat-Tg* T cells maintained the GVL effect. Donor T cells from *Cat-Tg* mice exhibited significantly lower inflammatory cytokine production and reduced donor T cell proliferation, while upregulating cytotoxic mediators that resulted in enhanced cytotoxicity. RNA sequencing data revealed changes in the expression of over 150 genes for CD4, and over 250 genes for CD8^+^T cells involved in essential aspects of immune response and GVHD pathophysiology. Transgenic over-expression of human β-catenin primarily affects CD8^+^ T cell phenotype. Altogether, our data suggest that β-catenin is a druggable target for developing therapeutic strategies to reduce GVHD while preserving the beneficial GVL effects following allo-HSCT treatment.

## Introduction

Allogeneic hematopoietic stem cell transplantation (allo-HSCT) is a curative treatment for patients with hematological malignancies, due to eradication of host malignant cells by donor T cells (graft-versus-leukemia or GVL) (1). In 40-70% of patients, the same donor T cells also attack healthy tissues like the gastrointestinal (GI) tract and liver, leading to graft-versus-host disease (GVHD) (2). The mortality rate due to GVHD is higher than 20% (3, 4). Therefore, therapeutic protocols that enhance the control of GVL and diminish the effects due to GVHD are essential for the treatment of leukemia. β-catenin signaling plays a critical role in T cell development and tissue homeostasis β-catenin pathways also play an important role in thymocyte development, β differentiation, polarization, and survival of mature T lymphocytes (6). Published data have β-catenin signaling is a key regulator of T cell development at various stages of β thymocyte differentiation (7). The transcription factor T Cell Factor-1 (*TCF-1*), encoded by the Tcf7 gene, and Lymphoid Enhancer Binding Factor-1 (LEF-1) are the downstream transcription effectors of the canonical Wnt signaling pathway (8). Both *Tcf7* and *Lef1* are highly expressed by naïve CD8^+^ T cells, and as these cells encounter antigen, the levels of *Tcf7* and *Lef1* decrease in these cells (9). There are several lines of evidence suggesting that Wnt/β-catenin signaling downregulates production of proinflammatory cytokines (10). These cytokines include IL-1β, IL-6, IL-8, and TNF-α. Wnt/β-catenin signaling is critical for T cell differentiation, effector functions, and migration (11). The activation of β-catenin signaling turns CD8^+^ T cells into Tfh-like cells (12). β-catenin signaling can also differentiate CD8^+^ T cells into effector cells (13).

There have been also reports that Wnt/β-catenin signaling pathways are highly expressed in naive CD8^+^ and memory CD8^+^ T cells, but expressed less in effector CD8^+^ T cells (11). Since the activation and maintenance of T cells are both required for anti-tumor immunity (GVL) and GVHD, we hypothesized that Wnt/β-catenin signaling may play an important role in these linked processes.

In this study, we used a novel mouse model expressing transgenic β mice,) that enhances the expression of the protein by 2-3 fold (8). We demonstrate that donor T cells from *Cat-Tg* mice do not induce GVHD in an MHC-mismatched mouse model, but still clear primary B-cell acute lymphoblastic leukemia (B-ALL) cells (14, 15). Our data also showed that *Cat-Tg* mice had fewer naïve CD8^+^ T cells and an increase in T cells with an activated phenotype that may trend towards exhaustion. Interestingly, our data show that recipient mice allogeneically transplanted with T cells from *Cat-Tg* mice had significantly decreased proinflammatory cytokines in serum, and the donor T cells showed lower levels of expansion. Unbiased analysis of gene expression using RNA sequencing showed that transgenic expression of β-catenin affected pathways like regulation of immune system processes, cell activation, lymphocyte activation, adaptive immune response, lymphocyte mediated immunity, MHC complex, cytokine production, TCR signaling pathways, T cell and B cell activation, T cell proliferation, Th1/Th2 cell differentiation, Th17 cell differentiation, signaling (NF-κβ, TNF, PLC-γ2, BLNK and others), GVHD, Allograft rejection, Autoimmunity, cell adhesion, and chemokine receptors in CD8^+^ and CD4^+^ T cells. We also observed that the similar pathways along with Hematopoietic cell lineage, GVHD, Allograft rejection, Th1, Th2, Th17 differentiation and other pathways was differentially regulated in CD8^+^ T by β-catenin over-expression. Importantly, trafficking of donor T cells to GVHD target organs is considered a hallmark of GVHD(16). Our data showed that transgenic expression of β-catenin specifically affects genes for cell chemotaxis, chemokine receptors, cell adhesion molecules, and cell migratory molecules. We confirmed the existence of a migration defect and examined tissue damage to target organs using histology. In summary, we provide a mechanistic understanding of the manner in which enhancement of the Wnt/β-catenin-TCF1/LEF1 pathway protects from GVHD while maintaining GVL. Thus, we have shown that β catenin and Wnt signaling are potential druggable targets to separate GVHD and GVT to improve allo-HSCT outcomes.

T cells from *Cat-Tg* mice have an activated phenotype. Both CD8^+^ and CD4^+^ T cells from *Cat-Tg* mice showed higher expression of CD44, and only in CD8^+^ T cells, higher expression of CD122 and Eomes, with no difference in T-bet expression. CD4^+^ T cells from *Cat-Tg* mice have significantly higher expression of PD-1, but no differences in CTLA-4 or TCF-1 expression.

CD8^+^ T cells from *Cat-Tg* mice also showed a trend towards increased expression of PD-1, but no difference in CTLA-4 or TCF-1 expression. Our data showed that there were less naïve and more central memory cells among CD8^+^ T cells from *Cat-Tg* mice. We observed that there were similar trends in CD4^+^ T cells from *Cat-Tg* mice, but these differences were not significant compared to CD4^+^ T cells from WT mice*-Tg* mice, but there were no significant differences compared to CD4^+^ T cells from WT mice.

We also found that T cells from *Cat-Tg* mice express more perforin and granzyme B than T cells from WT mice, showing that T cells from *Cat-Tg* mice have better cytotoxic function compared to T cells from WT mice. Next, we examined whether donor T cells from *Cat-Tg* mice had differences in serum cytokines post allo-HSCT. Our data show that recipient mice allogeneically transplanted with T cells from *Cat-Tg* mice had significantly decreased proinflammatory cytokines in the serum. On a cellular level, both CD8^+^ and CD4^+^ donor T cells from *Cat-Tg* mice expressed less IFN-γ, but had no difference in TNF-α compared to WT cells. We have also shown that both CD4^+^ and CD8^+^ T cells from *Cat-Tg* mice trend towards proliferating less than donor T cells from WT mice, but this effect was not statistically significant. Next, we examined TCR signaling and we did not observe any differences in PLC-γ1, ERK, or IRF-4 total protein level.

To investigate the transcriptomic changes from β atenin over-expression in T cells, we performed RNA sequencing as an unbiased approach to examine the mechanism behind the observed changes in T cell function and phenotype. Our data showed that on a transcriptomic level, pathways like regulation of immune system processes, defense responses, T cell and B cell activation, T cell proliferation, adaptive immune responses, immune system development, inflammatory responses, cytokine production, signaling (NF-κβ, TNF, PLC-γ2, BLNK and others), cell adhesion, and chemokine receptors in CD4^+^ T cells are significantly affected by β-catenin over-expression. We also observed that similar pathways, along with hematopoietic cell lineage, GVHD, allograft rejection, Th1, Th2, Th17 differentiation, and other pathways were differentially regulated in CD8^+^ T cells with β-catenin over-expression.

Trafficking of donor T cells to GVHD target organs is considered a hallmark of GVHD β-catenin specifically affects genes for cell chemotaxis, chemokine receptors, cell adhesion molecules, and cell migratory molecules. Finally, we have shown that CD8^+^ donor T cells from *Cat-Tg* mice have altered T cell proliferation. Both functional and genetic data show that T cells from *Cat-Tg* mice have differences in migration, chemokine receptor activity, chemotaxis and cell adhesion molecules of CD8^+^ T cells. Even though we saw significant differences in genes involved in these pathways in CD4^+^ T cells, the number of genes involved was less than in CD8^+^ T cells. We also did not observe any functional migration defect in CD4^+^ T cells.

For the first time, we have provided evidence that β-catenin has a significant impact on T cell functions in a disease model. This effect is due to changes in T cell phenotype, function, and gene expression. Thus, we have shown that β-catenin and Wnt signaling are potential druggable targets to separate GVHD and GVT to improve allo-HSCT outcomes.

## Materials and Methods

### Mice

*Cat-Tg* mice were described previously (18). C57BL/6, C57BL/6.SJL (B6-SJL), B6-Ly5.1 (B6.SJL-Ptprc^a^ Pepc^b^/BoyCrl) and BALB/c mice were purchased from Charles River or Jackson Laboratory. Mice aged 8-12 weeks were used, and all experiments were performed with age and sex-matched mice. Animal maintenance and experimentation were performed in accordance with the rules and guidance set by the institutional animal care and use committees at SUNY Upstate Medical University.

### Reagents, cell lines, flow cytometry

Monoclonal antibodies were purchased from Biolegend or eBioscience and were used at 1:100 dilution. Antibodies used included mouse anti-CD3(cat#100102), anti-CD28(cat# 102116), anti-CD3 BV605, anti-CD4-PE, anti-CD8-Pe/Cy7, anti-Eomes-Pe/Cy7, anti-CD44-Pacific Blue, anti CD122-APC, anti-CD62L-APC/Cy7, anti-T-bet-BV421, anti-CTLA4-PE, anti-PD1-BV785, anti-H-2K^b^-Pacific Blue, anti-TNF-α-FITC, anti-IFNγ-APC, anti-EdU-AF647, anti-CD45.1-FITC, anti-CD122-APC, anti-TCF-1-PE anti-CD45.2 APC. We performed multiplex ELISAs using the Biolegend LEGENDplex Assay Mouse Th Cytokine Panel kit (741043). D-Luciferin was purchased from Gold Bio (St Louis MO). Flow cytometry was performed on a BD LSR Fortessa (BD Biosciences). Data were analyzed with FlowJo software (Tree Star, Ashland, OR).

For cell sorting, T cells were purified with anti-CD90.2, or anti-CD4 and anti-CD8 magnetic beads using MACS columns (Miltenyi Biotec, Auburn, CA) prior to cell surface staining. FACS sorting was performed with a BD FACS Aria IIIu cell sorter (BD Biosciences). Cells were sorted into sorting media (50% FBS in RPMI) for maximum viability, or Trizol for RNAseq experiment. FACS-sorted populations were typically of > 95% purity. All cell culture reagents and chemicals were purchased from Invitrogen (Grand Island, NY) and Sigma-Aldrich (St. Louis, MO), unless otherwise specified. For signaling analysis, antibodies against PLCγ1, ERK, IRF-4, Granzyme, Perforin, and β ctin (total) were purchased from Cell Signaling Technology (Danvers, MA). The primary mouse B-ALL blasts cells (15) were transduced with luciferase, and cultured as described previously (19).

### Allo-HSCT and GVL studies

Lethally irradiated BALB/c mice (800 cGy, split into 2 doses of 400 cGy with 12 hours interval between) were injected intravenously with 10×10^6^ T cell-depleted bone marrow (_TCD_BM) cells with or without 1×10^6^ MACS purified CD3^+^ T cells. Donor T cells were taken from WT (C57Bl/6), WT Ly5.1 (B6.SJL-Ptprc^a^ Pepc^b^/BoyCrl), or *Cat-Tg* mice. For GVL experiments, *B*-cell acute lymphoblastic leukemia (B-ALL) primary blasts (14, 15) transduced with luciferase were cultured as described previously, and 1×10^5^ luciferase-expressing B-ALL blasts cells were used. Mice were evaluated once a week from the time of leukemia cell injection for more than 60 days post-transplant by bioluminescence imaging using the IVIS 200 Imaging System (Xenogen) (19). Clinical presentation of the mice was assessed 3 times per week by a scoring system that sums changes in 6 clinical parameters (for each parameter score was ranged from 0-2): posture, activity, fur texture, diarrhea, weight loss, and skin integrity (20). Mice were euthanized if they lost ≥ 30% of their initial body weight or became moribund.

### Cytokine production, cytotoxicity, and Edu incorporation assays

On Day 7 post-transplantation, serum from cardiac blood and single cell suspensions of splenocytes were obtained from allo-transplanted recipients. Serum IFN-γ, TNF-α, IL-5, IL-12, IL-6, IL-10, IL-9, IL-17A, IL-17F, IL-22, and IL-13 levels were determined by multiplex cytokine assays (Biolegend LEGENDplex)(15, 21). Splenocytes taken from allo-transplanted recipients were stimulated with anti-CD3/anti-CD28 (2.5ug/ml) for 6 hours in the presence of Golgiplug (BD Cytofix/Cytoperm Plus kit cat#555028) (1:1000). After incubation, the cells were fixed then permeabilized and stained intracellularly for cytokines (IFN-γ and TNF-α).

For the proliferation assay, recipient BALB/c mice were transplanted as described above (1×10^6^ CD3^+^ donor T cells and 10×10^6^ WT _TCD_BM), and recipient mice were injected at day 5-6 with 25mg/kg EdU (20518 from Cayman Chemicals) in PBS. At day 7, the recipient mice were euthanized and lymphocytes from spleen were obtained. Cells were processed and stained using an EdU click chemistry kit (C10424 from Invitrogen), and also stained for H2Kb, CD3, CD4, and CD8 to identify donor cells as perversely described (22).

For cytotoxicity assays, luciferase-expressing B-ALL cells were seeded in 96-well flat bottom plates at a concentration of 3×10^5^ cells/ml. D-firefly luciferin potassium salt (75 μg/ml; Caliper Hopkinton, MA) was added to each well and bioluminescence was measured with the IVIS-50 Imaging System. Subsequently, effector cells (MACS-purified) were added at 40:1, 20:1, and 10:1 effector-to-target (E:T) ratios and incubated at 37°C for 4 hours. Bioluminescence in relative luciferase units (RLU) was then measured for 1 minute. Cells treated with RIPA lysis buffer was used as a measure of maximal killing. Target cells incubated without effector cells were used to measure spontaneous death. Triplicate wells were averaged and percent lysis was calculated from the data using the following equation: % specific lysis = 100X (spontaneous death RLU–test RLU)/(spontaneous death RLU– maximal killing RLU) (23)

### Migration assays

Lethally irradiated BALB/c mice were injected intravenously with 10×10^6^ WT T cell-depleted bone marrow (_TCD_BM), and a 1:1 mixture of WT (B6-Ly5.1) CD45.1^+^ MACS-purified CD8^+^ and CD4^+^ T cells (checked for 1:1 ratio with flow) with either WT (C57BL/6) CD45.2^+^ cells or *Cat-Tg* CD45.2^+^ cells (total 1×10^6^ cells). The donor cells were checked pre-transplant for a 1:1 ratio of donor types, and a 1:1 ratio of CD4:CD8 T cells within each donor type. Seven days post-transplantation, the mice were sacrificed and lymphocytes from the liver, small intestine, and spleen were isolated. Livers were perfused with PBS, dissociated, and filtered with a 70μm filter. The small intestines were washed in media, shaken in strip buffer at 37°C for 30 minutes to remove the epithelial cells, and then washed, before digesting with collagenase D (100 mg/ml) and DNase (1mg/ml) for 30 minutes in 37°C, and followed by filtering with a 70 μm filter.

Lymphocytes from the liver and intestines were further enriched using a 40% Percoll gradient. The cells were analyzed for H2K^b^, CD45.1^+^ and CD45.2^+^, CD3^+^, CD8^+^ and CD4^+^ by flow cytometry as described before (15, 21).

### RNA sequencing

3 Recipient BALB/c mice for each group were short-term transplanted as described above (1×10^6^ CD3^+^ donor T cells and 10×10^6^ WT _TCD_BM), and at day 7, recipient mice were euthanized and splenocytes were obtained for post-transplant samples. 3 WT or *Cat-Tg* mice also were euthanized and fresh splenocytes were isolated for pre-transplanted samples. CD4^+^ and CD8^+^ T cells from each pre- and post-transplanted mouse were FACS sorted as described above. These cells were all sorted into Trizol and brought to the Molecular Analysis Core (SUNY Upstate) https://www.upstate.edu/research/facilities/molecular-analysis.php for RNA extraction and library prep, followed by RNA sequencing analysis at the University at Buffalo Genomics Core http://ubnextgencore.buffalo.edu. We generated RNA sequencing data from four groups for each cell subset (CD4/CD8): WT-pre tx and *Cat-Tg* pre tx cells (prior to transplantation); and WT-Day7 tx, *Cat-Tg* Day-7 tx (7 days post-transplantation). We were unable to sort enough donor T cells from small intestines and liver of the recipient mice that received *Cat-Tg* CD3^+^ T cells. All data were processed and analyzed using the R programming language (Version 4.0.4), the RStudio interface (Version 1.4.1106), and Bioconductor. For pseudoalignment and gene expression transcript abundance of samples was computed by pseudoalignment with Kallisto_41_(version 0.46.2). Transcript per million (TPM) values were then normalized and fitted to a linear model by empirical Bayes method with the Voom and Limma R packages (24, 25) and differential gene expression was defined as a Benjemini and Hochberg adjusted *p*-value-False L 0.1. For functional enrichment analysis, the g: Profiler (26) toolset, g:GOSt tool was used to illustrate Manhattan plot. Gene Set Enrichment Analysis (GSEA) was performed against the Molecular Signatures Database (MSigDB)(27) using the Hallmark pathways collection. Data will be deposited (https://www.ncbi.nlm.nih.gov/geo/). The RNAseq experiment described here was performed as part of the experiment described in other recent publications from our lab (15, 28). Therefore, the data generated for WT pre- and post-transplanted samples (CD4 and CD8) is the same as that shown in the papers mentioned, but here these data are compared to data for Cat-Tg mice.

### Western blotting

Cells were lysed in freshly prepared lysis buffer (RIPA buffer (Fisher Scientific cat#PI89900) + cOmplete protease inhibitor cocktail (Sigma Aldrich cat#11697498001) and centrifuged at 14000 rpm for 10 minutes at 4°C. Aliquots containing 1×10^6^ cells were separated on 12-18% denaturing polyacrylamide gel and transferred to nitrocellulose membranes for immunoblot analysis using specific Abs.

### Histopathological Evaluation

Lethally irradiated recipient mice were transplanted with 10×10^6^ T cell-depleted bone marrow cells, and 1×10^6^ CD3^+^T cells from WT or *Cat-Tg* mice. At day 7 post-transplantation, recipient mouse livers and smalls intestine were obtained and fixed in 10% neutral buffered formalin, then were sectioned and stained with H&E by the Histology Core at Cornell University (https://www.vet.cornell.edu/animal-health-diagnostic-center/laboratories/anatomic-pathology/services). Obtained tissues were graded for GVHD by a pathologist (A.M), who was blinded to the study group and disease status. Links for grading criteria: http://surgpathcriteria.stanford.edu/transplant/skinacutegvhd/printable.html, http://surgpathcriteria.stanford.edu/transplant/giacutegvhd/printable.html, http://surgpathcriteria.stanford.edu/transplant/livergvhd/printable.html. Statistical analysis was performed using Mann-Whitney U test.

### Statistics

All numerical data are reported as means with standard deviation unless otherwise noted in figure legends. Data were analyzed for significance with GraphPad Prism v7. Differences were determined using one-way or two-way ANOVA and Tukey’s multiple comparisons tests, chi-square test or with a student’s t-test when necessary. We used Mann Whitney U test for analysis of GVHD grades. P-values less than or equal to 0.05 are considered significant. All transplant experiments are done with N=3 mice per group, and repeated at least twice, according to power analyses unless otherwise specified. Mice are sex-matched, and age-matched as closely as possible.

## Results

### Donor T cells from *Cat-Tg* mice do not induce GVHD but maintain GVL function

To determine whether mouse over-expression of β-catenin signaling on donor CD4^+^ and CD8^+^ T cells in β an allotransplant model, using C57Bl/6 background mice (MHC haplotype^b^) as donors and BALB/c mice (MHC haplotype ^d^) as recipients (15). To induce GVHD, we used MHC-mismatched donors and recipients. T cell-depleted bone marrow cells from WT mice, and T cells from C57BL/6 (B6) WT or *Cat-Tg* mice were injected into irradiated BALB/c mice along with luciferase-expressing B-cell acute lymphoblastic leukemia (B-ALL-*luc)* tumor cells (14, 15).

Lethally irradiated BALB/c mice were injected intravenously with 10×10^6^ wild-type (WT) Tcell-depleted donor BM cells along with 1×10^6^ MACS-sorted donor CD3^+^ T cells along with 1×10^5^ B-ALL-*luc* blast cells as described (14, 15, 21). Recipient BALB/c mice were monitored for cancer cell growth using IVIS bioluminescence imaging for over 60 days (**Fig. 1A**)(**15, 21**). While leukemia cell growth was observed in mice given bone marrow without T cells, leukemia cell growth was not seen in mice transplanted with T cells from either WT or *Cat-Tg* mice. As expected, mice transplanted with WT T cells cleared the leukemia cells (**Fig. 1A**) but suffered significantly from GVHD (**Fig 1B-D**). In contrast, mice transplanted with *Cat-Tg* T cells cleared the leukemia cells (**Fig. 1A**) and displayed minimal signs of GVHD (**Fig. 1B-D**). All animals transplanted with *Cat-Tg* T cells survived for more than 65 days post-allo-HSCT (**Fig. 1B**), with significantly reduced weight loss and clinical scores compared to those transplanted with WT Tcells (scored based on weight loss, posture, activity, fur texture, and skin integrity, and diarrhea as previously described) (20), (**Fig. 1C-D**). Quantification of tumor bioluminescence showed that mice given WT or *Cat-Tg* T cells cleared the tumor cells, while tumor burden remained high for mice only given bone marrow (**Fig. 1E**). Our results indicate that donor T cells from *Cat-Tg* mice are dispensable for anti-leukemia immunity, but required for GVHD damage.

**Figure 1.**
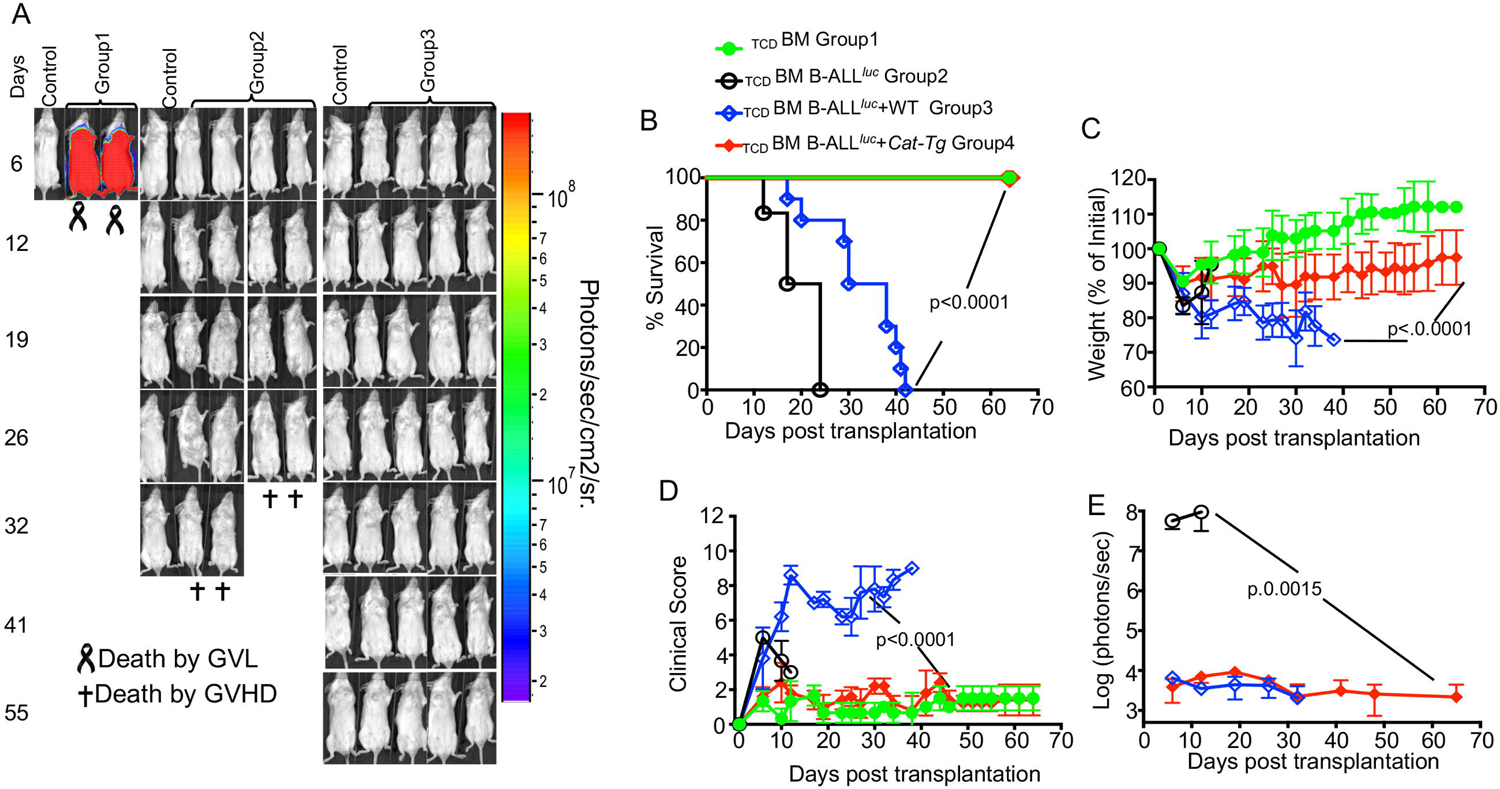
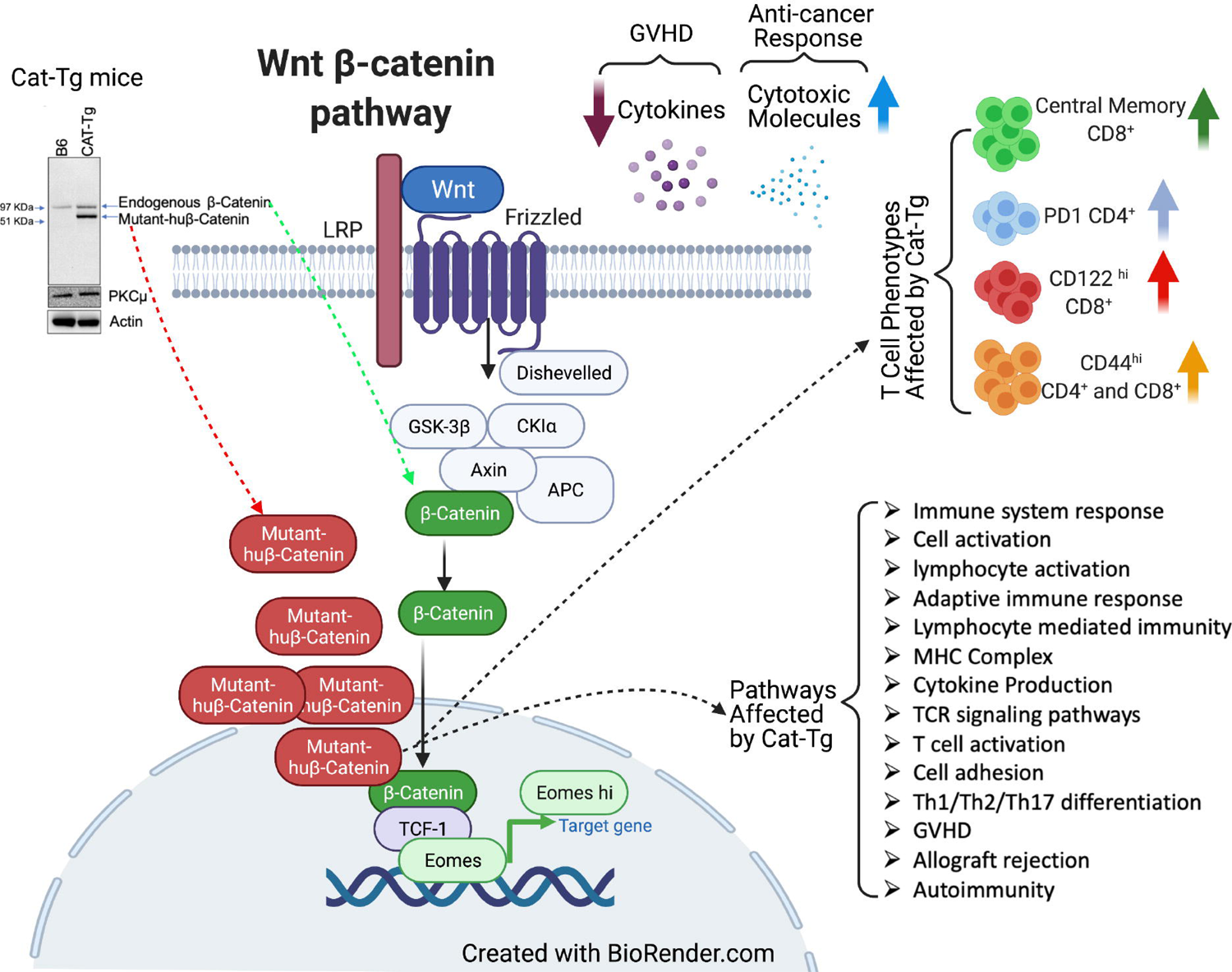
β-catenin over-expression retains the GVL effect but avoids GVHD during allo-HSCT. 1×10^6^ purified WT or *Cat-Tg* CD3+ T cells were transplanted into lethally irradiated BALB/c recipient mice along, with 10×10^6^ T cell depleted bone marrow cells from WT mice and 1×10^5^ B-ALL-*luc* cells. Host BALB/c mice were imaged using the IVIS50 system 3 times a week. Group 1 received T cell-depleted bone marrow only (labeled as _TCD_BM). Group 2 received 10×10^6^ _TCD_BM from WT mice and 1×10^5^ B-ALL-*luc* cells (_TCD_BM+B-ALL *^luc^*^+^). Group 3 was transplanted with 10×10^6^ _TCD_BM from WT mice and 1×10^6^ purified WT CD3^+^ T cells, along with 1×10^5^ B-ALL*^luc^*^+^ cells (_TCD_BM+B-ALL*^luc^*^+^ WT). Group 4 received 10×10^6^ _TCD_BM from WT mice and 1×10^6^ purified *Cat-Tg* CD3^+^ T cells, along with 1×10^5^ B-ALL-*luc*+ cells (_TCD_BM+B-ALL*^luc^*+ *Cat-Tg*). **(A)** Recipient BALB/c mice were imaged using IVIS50 3 times a week. The mice were also monitored for **(B)** survival, **(C)** changes in body weight, and **(D)** clinical score for 65 days post BMT. (**E**) Quantitated luciferase bioluminescence of tumor growth. Statistical analysis for survival and clinical score was performed using log-rank test and one-way ANOVA, respectively. For weight changes and clinical score, one representative of 2 independent experiments is shown (n = 3 mice/group for BM alone; n = 5 experimental mice/group for all three other groups) Survival is a combination of 2 experiments. *Note: Control mouse is a naïve mouse used as a negative control for BLI*.

### β-catenin affects T cell phenotype and cytotoxic function

β-catenin affects T cell phenotype, we MACS-purified T cells from *Cat-Tg* WT mice by CD90.2-positive selection. We examined the effects of β-catenin over-expression on CD4^+^ and CD8^+^ T cells in comparison to T cells from WT C57Bl/6 mice. Our data showed that CD8^+^ T cells from *Cat-Tg* mice exhibit an innate memory phenotype (IMP) (15, 29), as indicated by expression of high levels of CD44, CD122, and a key transcription factor Eomesodermin (Eomes). We did not observe any changes in T-bet expression. CD8^+^ T cells from *Cat-Tg* mice showed a trend towards increased levels of PD-1, but had no differences in CTLA-4 expression compared to CD8^+^ T cells WT mice. We also did not observe significant differences in TCF-1 expression (**Fig. 2A-D**). Next, we examined the β-catenin on CD4^+^ T cells. Our data show that CD4^+^ T cells from *Cat-Tg* mice also express higher levels of CD44 but had no differences in CD122, Eomes, or T-bet expression compared to CD4^+^ T cells from WT mice. CD4^+^ T cells from *Cat-Tg* mice also express significantly higher percentage of PD-1 but have no differences in CTLA-4 or TCF-1 compared to CD4^+^ T cells from WT mice (**Fig. 2C-D**). Our data shows that CD4^+^ T cells from *Cat-Tg* mice were no difference compared to CD4^+^ from WT mice. observed in effector memory or transitional to activation state (**Fig.2** E) (**Supp. Fig.1A**). Next, we examined whether CD8^+^ T cells from *Cat-Tg* mice have changes in memory subsets. We observed a significant decrease in naïve CD8+ T cells, no significant differences in the transitioning/activating cells, but significantly increased central memory CD8^+^ T cells. No significant differences in effector memory CD8^+^ cells from *Cat-Tg* mice compared to CD8^+^ T cells from WT mice was observed (**Fig. 2F**) (**Supp. Fig.1B**).

**Figure 2.**
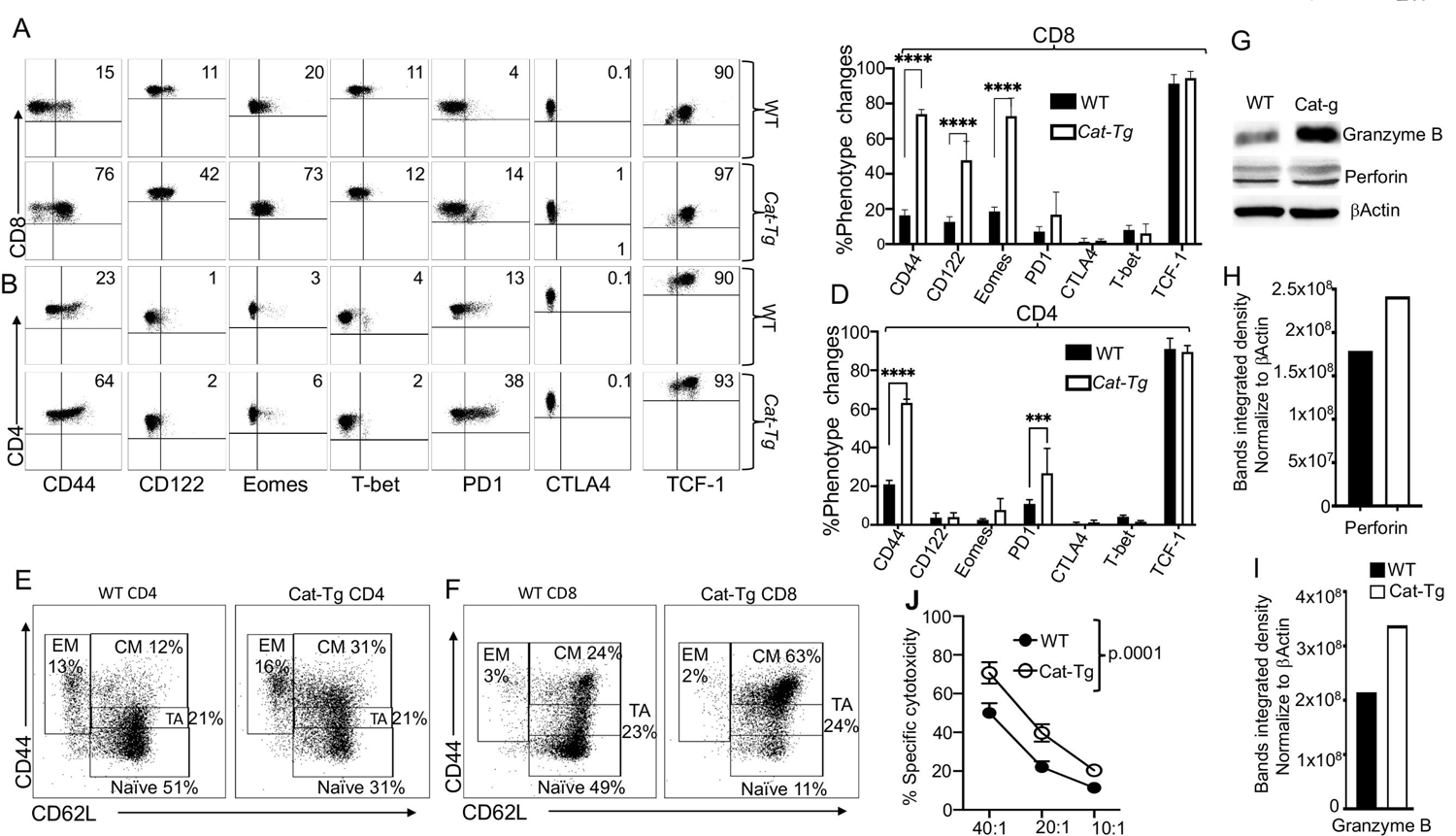
T cells from Cat-Tg mice exhibit enhanced T cell IMP phenotypes and enhanced GVL cytotoxicity. (A-B) Purified CD3^+^ T cells from WT and *Cat-Tg* T cells were examined for expression of CD44, CD122, Eomes, T-bet, PD-1, CTLA-4 and TCF-1 by flow cytometry. These markers were examined for both **(A)** CD8^+^ and **(B)** CD4^+^ T cells. **(C-D)** Quantification of 3 different experiments of CD44, CD122, Eomes, T-bet, PD-1, CTLA-4 and TCF-1 in **(C)** CD8^+^ or in **(D)** CD4^+^. (**E)** CD8^+^ **(F)** CD4^+^ T cells from WT and *Cat-Tg* mice were examined for effector memory, central memory, transitioning to activation (TA), and naïve population frequencies. **(G)** Purified T cells were examined for expression of perforin, granzyme B, and β-actin by western blot. **(H-I)** Quantitative analysis of perforin **(H)** and granzyme B **(I)** expression from western blot data, normalized against β−Actin. **(J)** *Ex vivo* purified T cells were used in a cytotoxicity assay against primary tumor target B-ALL*luc*+ cells at a 40:1, 20:1, or 10:1 effector to target ratio. Statistical analysis was performed using two-way ANOVA, one-way ANOVA confirmed by Student’s *t-*test, p-values are presented. Symbol meanings for P-values are: ns - P > 0.05; * - P ≤ 0.05; ** - P ≤ 0.01; *** - P ≤ 0.001; **** - P ≤ 0.0001(n = 3 mice per group)

To examine whether T cells from *Cat-Tg mice* have cytotoxic function, we purified CD3^+^ T cells from *Cat-Tg* mice and WT mice, and performed a western blot on the cell lysates. Our western data show that CD3^+^ T cells from *Cat-Tg* mice express significantly higher levels of granzyme B and perforin than CD3^+^ T cells from WT mice (**Fig. 2G-I**). We quantified the band integrated density and normalized it to β−actin using Image Lab software (**Fig. 2H-I**). Next, we examined whether CD8^+^ T cells from *Cat-Tg* mice could mount a cytotoxic response, using a cytotoxicity assay against primary B-ALL cells in different effector to target ratios (14, 15, 21). We found that CD8^+^ T cells from *Cat-Tg* mice effectively killed significantly more primary leukemia cells *in vitro* than CD8^+^ T cells from WT mice (**Fig. 2J**). Our findings demonstrate that CD8^+^ T cells from *Cat-Tg* mice have enhanced activation markers, significantly altered CD8^+^ T cells phenotype, enhanced expression of Granzyme B and Perforin, and exert better cytotoxicity against primary leukemia cells than CD8^+^ T cells from WT mice.

### Wnt/β catenin over-expression results in reduced cytokine production and donor T cell proliferation without affecting TCR signaling

The conditioning regimen for allo-HSCT elicits an increase in the production of inflammatory cytokines by donor T cells, known as a “cytokine storm”(30, 31). This is considered one of the hallmarks of GVHD pathogenesis (32). We assessed cytokine production by *Cat-Tg* T cells in our allo-HSCT model (B6 BALB/c) by examining the levels of serum inflammatory cytokines. We observed that recipient BALB/c mice treated with CD3^+^ T cells from *Cat-Tg* mice expressed significantly less IFN-γ, TNF-α, IL-5, IL-2, IL-6, IL-10 and IL-22 in serum compared to recipient BALB/c treated with CD3^+^ T cells from WT mice (**Fig. 3A**). We did not observe differences in IL-4, IL-9, IL-17A, IL-17F, and IL-13 on day 7 post allotransplantation (**Fig. 3A**). We also examined donor CD8^+^ or CD4^+^ T cells from secondary lymphoid organs of recipients using anti-H2K^b^ antibodies (H2K^b^ is expressed by donor C57Bl/6 cells). Ex *vivo* donor T cells were cultured for 6 hours with GolgiPlug and stimulated with anti-CD3/CD28 antibodies (**Fig. 3B-C**) or left unstimulated, followed by analysis of IFN-γ and TNF-α cytokine production. *Cat-Tg* CD8^+^ or CD4^+^ T cells produced significantly less inflammatory IFN-γ when stimulated via anti CD3/CD28 antibodies, but we did not observe any differences in TNF-α expression (**Fig. 3B-E**).

**Figure 3.**
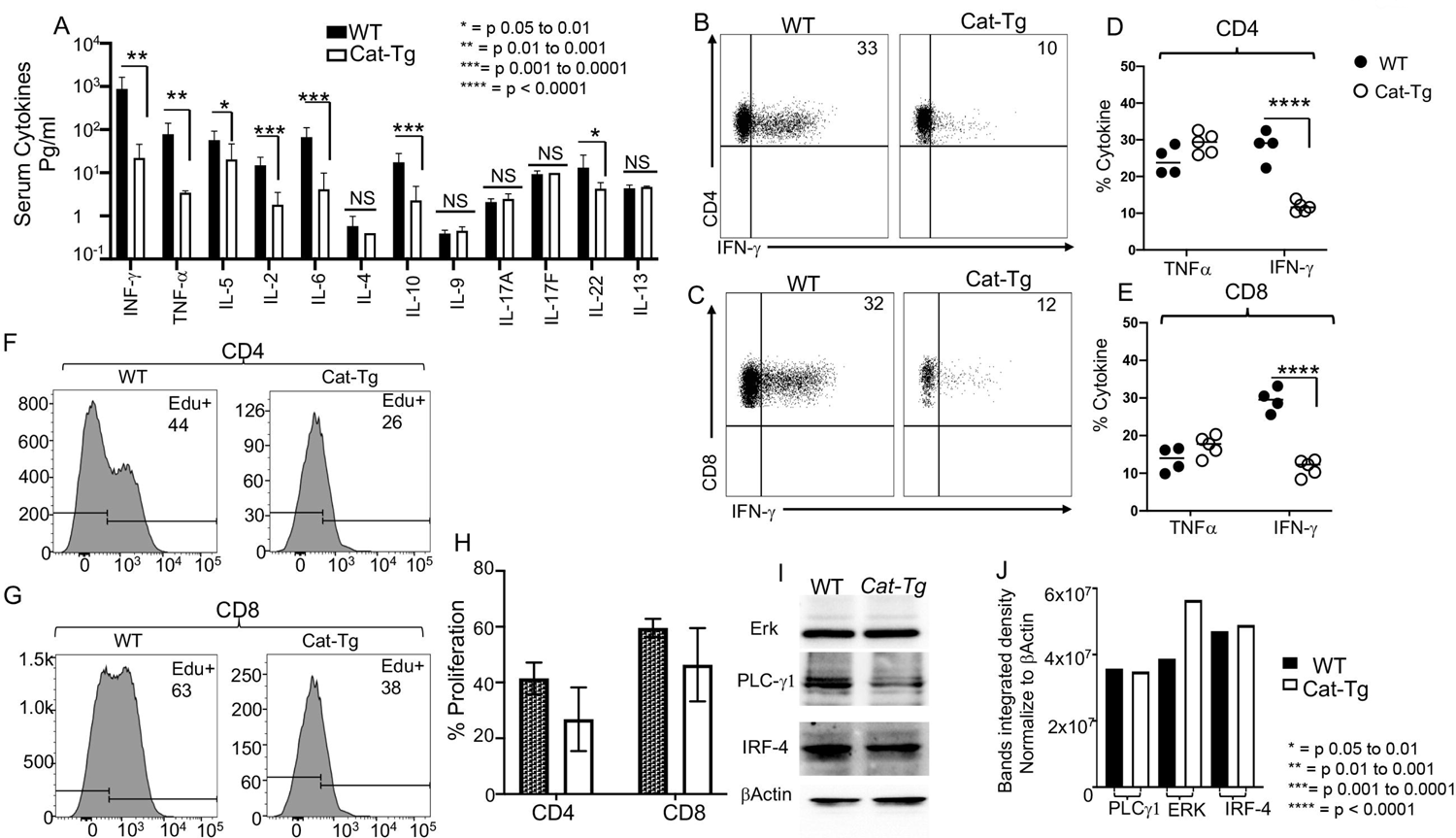
Over-expression of β-Catenin reduces T cell inflammatory cytokine production and proliferation without affecting signaling molecules. **(A)** 1×10^6^ purified WT or *Cat-Tg* CD3^+^ T cells were transplanted with _TCD_BM into irradiated BALB/c mice. At day 7 post allo-HSCT, recipient BALB/c were euthanized and serum cytokines (IFN-γ, TNF-α, IL-2, IL-5, IL-6, IL-4, IL-10, IL-9, IL17A, IL-17F, IL-22, and IL-13) were determined by multiplex ELISA. **(B-C)** Intracellular IFN-γ and TNF-α expression by donor CD4 **(B**) and CD8 **(C)** T cells after 6 hr stimulation with anti-CD3/anti-CD28 and GolgiPlug, as determined by flow cytometry. **(D-E)** Quantified IFN-γ and TNF-α expression for **(B-C)**. (F-H) *Ex vivo* proliferation of donor CD4^+^ or CD8^+^ T cells from *Cat-Tg or* WT mice. Lethally irradiated recipient BALB/c mice were transplanted as mentioned above, with either WT or *Cat-Tg* donor CD3^+^ T cells. Recipient mice were given EdU in PBS i.p. (25 mg/kg in 100μl) for last 2 days. At 7 days post-allotransplantation, recipient mice were sacrificed and examined for proliferation by EdU incorporation via flow cytometry. **(I)** Purified CD3^+^ WT and *Cat-Tg* T cells were examined for expression of ERK, PLCγ −1, and IRF-4 by western blot. **(J)** Quantitative analysis from western blots in **(I)**, using ImageLab software to normalize to β−Actin. Symbol meanings for P values are: ns - P > 0.05; * - P ≤ 0.05; ** - P ≤ 0.01; *** - P ≤ 0.001; **** - P ≤ 0.0001. (For Fig 3A-G, n = 5 mice per group, for Fig 3 H-I, n=3 mice used per group)

We next examined donor T cell proliferation using an EdU incorporation assay. We utilized short term allo-transplantation as described above, and recipient were injected with EdU in PBS on day 5 and 6 post-transplant. Seven days post allo-transplantation, splenocytes were obtained from recipients, and donor cells (identified by H2K^b+^, CD3^+^ and CD4^+^ or CD8^+^) were examined for proliferation by EdU incorporation. Both donor CD4^+^ and CD8^+^ T cells from *Cat-Tg* mice showed a trend toward reduced proliferation compared to donor T cells from WT mice, but this effect was not significant (**Fig. 3F-H**).

We have recently shown that modulating TCR signaling through ITK causes T cells to acquire an innate-like memory phenotype (IMP), distinguished by higher expression of CD44, CD122 and Eomes due attenuated TCR signaling (15, 21, 29), Modulation of ITK also affects ERK, PLCγ-1 and IRF-4 expression level(15). We did not observe significant differences in any of these signaling molecules on T cells from *Cat-Tg* or WT mice (**Fig. 3I-J**). Our data suggest that T cells from *Cat-Tg* mice leads to reduced inflammatory cytokine production and reduced proliferation of allogeneically transplanted T cells in a major mismatch model. Our data also β-catenin does not attenuate TCR signaling. These findings β support our observations that GVHD severity is reduced by over-expression of β-catenin.

### Wnt/β catenin expression regulates gene expression in T cells during GVHD

As an unbiased approach to further explore differences between CD4^+^ or CD8^+^ T cells from WT and *Cat-Tg* mice, we employed RNA sequencing analysis. We examined the differences in gene expression between WT and *Cat-Tg* CD4^+^ and CD8^+^ T cells before and following allo-HSCT. We sort-purified donor WT and *Cat-Tg* CD4^+^ or CD8^+^ T cells from freshly isolated splenocytes, and called these pre-transplanted cells (pre-tx). We also MACS purified CD3^+^ T cells from WT or *Cat-Tg* mice and transplanted them along with T cell-depleted bone marrow cells into the tail vein of lethally irradiated BALB/c recipients. At day 7 post-transplantation, we sort-purified donor WT and *Cat-Tg* CD4^+^ or CD8^+^ T cells (using H-2K^b^ antigen expressed by donor T cells) from recipients, and called these post-transplant day 7 samples (Day 7-tx). The cells were sorted into Trizol reagent and transcriptionally profiled. Principal component analysis (PCA) of CD4^+^ T cells identified four clusters of samples, which clearly separated the pre-transplanted WT or *Cat-Tg* and post-transplanted WT or *Cat-Tg* populations along PC1 (39.5%) and PC2 (17%) (**Fig. 4A**). Further analysis of these cell populations identified ∼150 differentially expressed genes (DEGs; FDR <0.1) between WT and *Cat-Tg* (**Supplementary Table 1**), of which 74 genes were downregulated and 76 genes were upregulated. These genes were plotted on a volcano plot (**Fig. 4B**). The use of a Spearman correlation method associated with hierarchical clustering of CD4^+^ T cell samples showed that the WT and *Cat-Tg* post-transplant samples were most similar (while still clustering separately). Cat-Tg pre-transplant samples were also more similar to post-transplant samples than the WT pre-transplant samples were (**Fig 4C**). DEGs between WT and *Cat-Tg* in CD4^+^ T cells were averaged by group and gene co-regulation was determined by hierarchical clustering, using Pearson correlation with a grouping cutoff (*k*) of 4. Each generated module contributed to the different pathway enrichment. Gene expression is averaged by group 3) for clarity and displayed as *z* score across each row (**Fig. 4C**). Functional pathway L analysis of these genes revealed that these DEGs in CD4^+^ T cells are involved in numerous biological pathways including immune system process, defense response, cell activation, adaptive immune response, cytokine production, regulation of T cell activation and proliferation, MHC protein complex binding, regulation of programmed cell death, chemokine regulation, autoimmunity, signaling like TNF, NF-kappa B and others which are illustrated in the Manhattan plot. The relevant pathways related to our project were highlighted and ID details are given about each pathway. This information is shown in the table according to the ID of each pathway (**Fig. 4D**). The whole list of differentially expressed functional pathways involved in CD4^+^ T cells are listed in **Supplementary Table 2**. When we performed GSEA analysis of these genes using the Hallmark pathways collection from Molecular Signatures Database (MSigDB)(27) we observed that TNF-α signaling via NF-kβ is enriched in WT compared to *Cat-Tg* in pre-transplanted samples (**Sup. Fig.2A**). Interestingly, in post-transplant day 7 samples, TNF-α signaling via NF-kβ, IFN-γ, IFN-α, inflammatory responses, and apoptosis pathways are enriched in *Cat-Tg* rather than WT (**Sup. Fig.2B**).

**Figure 4.**
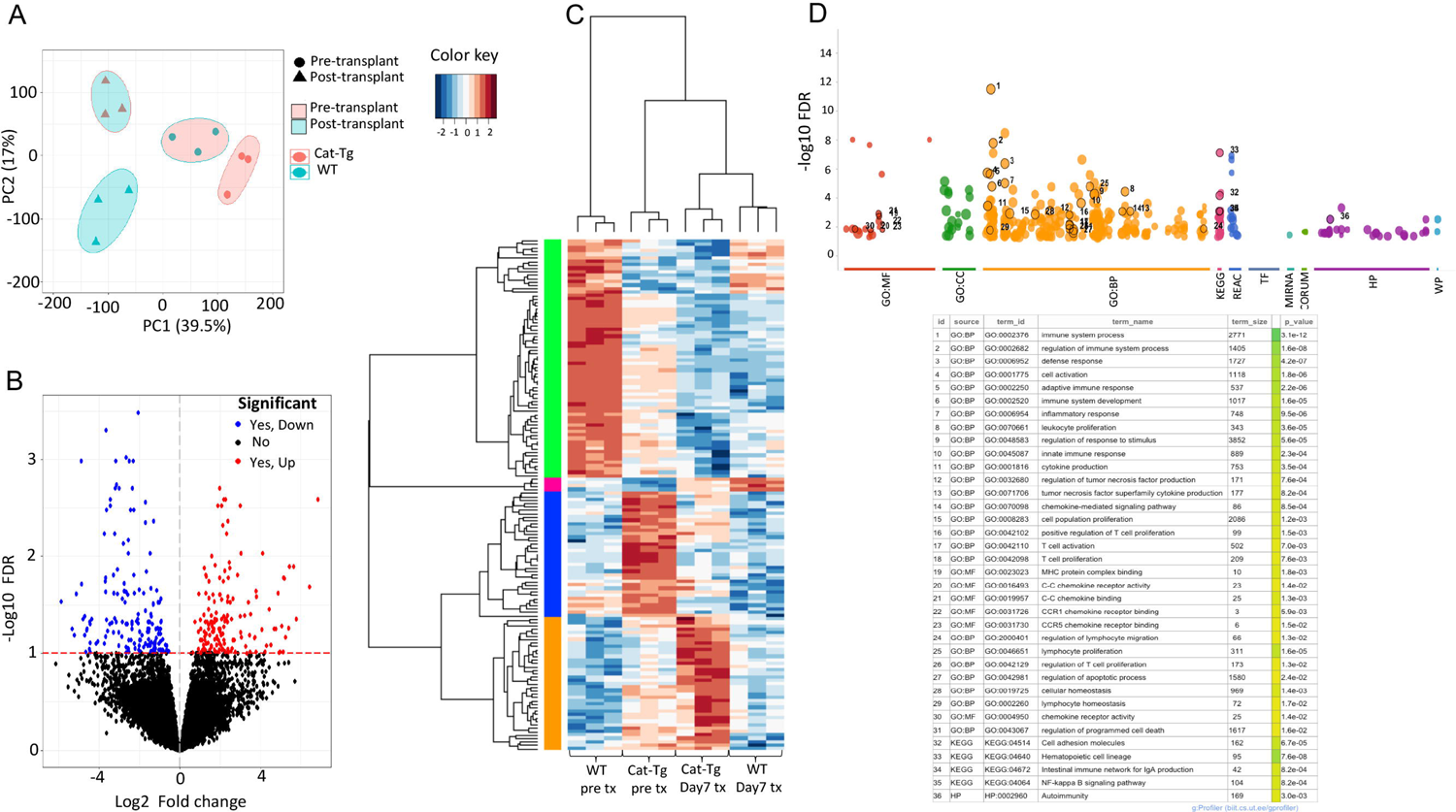
β-catenin over-expression differentially regulates gene expression in CD4^+^ T cells during GVHD. **(A)** PCA analysis showing clustering of pre-transplanted and post-transplanted day 7 CD4^+^ T cells by strain and by timepoint. Samples grouped for strain by color and for timepoint by shape. All replicates are shown (n = 3). **(B)** Volcano plot displaying differentially expressed genes (FDR<0.1) between *Cat-Tg* and WT day 7 CD4^+^ T cell samples. Genes up-regulated (red) and down-regulated (blue) are labeled. **(C)** Hierarchical clustering and heat map illustrating expression of genes compared between different groups selected by strain and timepoint. All replicates are shown (n = 3) for each group. **(D)** Manhattan plot showing functional pathway analysis of CD4^+^ T cells. Using the online tool gProfiler and the ordered g:GOSt query, we assessed which biological processes (BP) were linked to the list of 150 significantly differentially expressed genes from CD4^+^ T cells. The x-axis represents functional terms that are grouped and color-coded by data sources [molecular function (MF), biological process (BP), cell component (CC)]. The y-axis shows the adjusted enrichment p-values on a negative log10 scale. Adjusted p-values g:GOSt used Bonferroni correction and a threshold of 0.1. On the table, adjusted p-values were color coded as yellow for insignificant findings to dark blue with highest significance. (n = 3 mice per group)

We also analyzed the CD8^+^ T cell samples and performed principal component analysis (PCA) of CD8^+^ T cells. We again identified four groups in which pre-transplanted samples were clearly separated, while post-transplant samples were not separated as well as pre-transplant samples along PC1 (32.2%) and PC2 (15.3%) (**Fig. 5A**). Further analysis of these cell populations identified ∼250 differentially expressed genes (DEGs; FDR <0.1) between WT and *Cat-Tg* (**Supplementary Table 3**), of which 175 genes were downregulated while 80 genes were upregulated, and these DEGs are plotted on a volcano plot (**Fig. 5B**). The use of a Spearman correlation method associated with hierarchical clustering of CD8^+^ T cell samples showed the same clustering as of CD4^+^ T cell samples. (**Fig. 5C**). DEGs between *Cat-Tg* and WT in CD8^+^ T cell were averaged by group and gene co-regulation was determined by hierarchical clustering using Pearson correlation with a grouping cutoff (*k*) of 4. Each generated module contributed to L 3) for clarity and displayed as *z* score across each row. (**Fig. 5C**).

**Figure 5.**
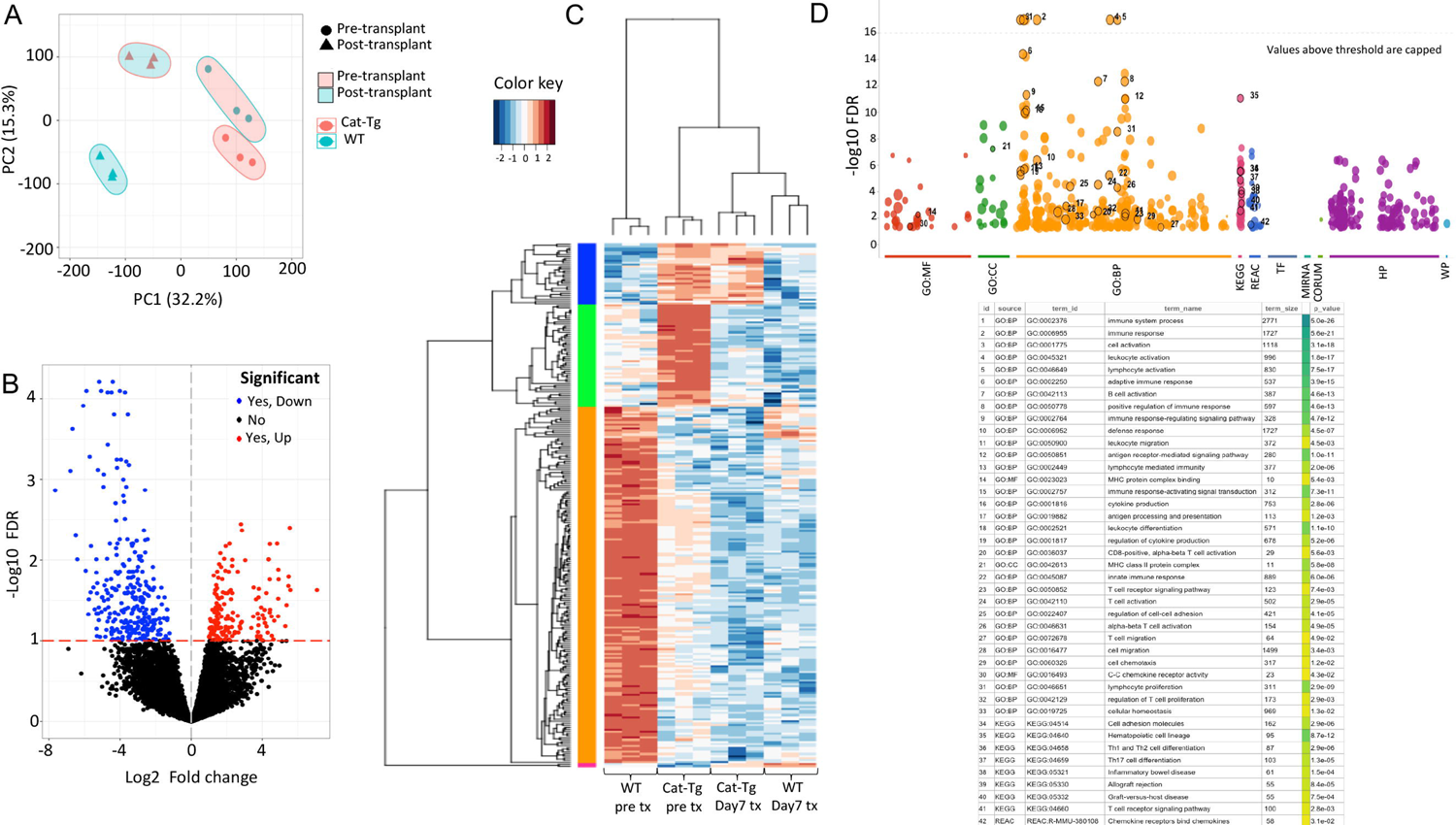
β catenin over-expression differentially regulates gene expression in CD8 T cells during GVHD. **(A)** PCA analysis showing clustering of pre-transplanted and post-transplanted day 7 CD8^+^ T cells by strain and by timepoint. Samples grouped for strain by color and for timepoint by shape. All replicates are shown (n = 3). **(B)** Volcano plot displaying differentially expressed genes (FDR<0.1) between *Cat-Tg* and WT day 7 CD8^+^ T cell samples. Genes up-regulated (red) and down-regulated (blue) are labeled. **(C)** Hierarchical clustering and heat map illustrating expression of genes compared between different groups selected by strain and timepoint. All replicates are shown (n = 3) for each group. **(D)** Manhattan plot showing functional pathway analysis of CD8^+^ T cells. Using the online tool gProfiler and the ordered g:GOSt query, we assessed which biological processes (BP) were linked to the list of 150 significantly differentially expressed genes from CD8^+^ T cells. The x-axis represents functional terms that are grouped and color-coded by data sources [molecular function (MF), biological process (BP), cell component (CC)]. The y-axis shows the adjusted enrichment p-values on a negative log10 scale. Adjusted p-values g:GOSt used Bonferroni correction and a threshold of 0.1. On the table, adjusted p-values were color coded as yellow for insignificant findings to dark blue with highest significance. (n = 3 mice per group)

Functional pathway analysis revealed that the DEGs in CD8^+^ T cells are involved in similar biological pathways to the DEGs seen in CD4^+^ T cell, as well as in TCR signaling, hematopoietic cell lineage, Th1, Th2 and Th17 cell differentiation, inflammatory bowel disease, allograft rejection, graft-versus-host-disease, and others which are illustrated in the Manhattan plot. The important pathways related to our project were highlighted and ID details about each pathway are shown in the table. This information is given according to the ID of each pathway (**Fig. 5D**). The whole list of differentially expressed functional pathways involved in CD8^+^ T cells is listed in **Supplementary Table 4**. When we performed GSEA analysis of these genes using the Hallmark pathways collection from Molecular Signatures Database (MSigDB)(27), we observed that TNF-α signaling via NF-kβ, κRAS signaling, IFN-γ response, inflammatory response, IL6-JAK-STAT3 signaling, IL2-STAT-5 signaling, and allograft rejection pathways are enriched in WT compared to *Cat-Tg* in pre-transplanted samples **(Sup. Fig.3A)**. Once again, in post-transplant day 7 samples, TNF-α signaling via NF-kβ, κRAS signaling, PI3K/AKT/MTOR signaling, IFN-γ and IFN-α response, inflammatory response, IL6/JAK/STAT3 signaling, IL2/STAT-5 signaling, and allograft rejection pathways are enriched in *Cat-Tg* compared to WT **(Sup. Fig.3B)**. Altogether these data suggest that β catenin plays an important role in regulating the immune response to allo-antigens by controlling gene expression programs in mature and alloactivated T cells.

### β-catenin over-expression specifically affects CD8^+^ T cell functions

The pathogenesis of GVHD involves migration of donor T cells into the target organs in the recipient, including liver, small intestine, and skin (33, 34). GVHD occurs in a subset of organs and involves early migration of alloreactive T cells into these organs followed by T cell expansion and later tissue destruction(16, 35). To examine whether over-expression of Wnt/β catenin affects donor T cell migration, irradiated BALB/c recipient mice were injected with CD8^+^ T cells and CD4^+^ T cells from *Cat-Tg* (CD45.2^+^) and WT B6-Ly5.1 (CD45.1^+^) mice mixed at a 1:1 ratio of WT: *Cat-Tg* (**Fig. 6A**). A total of 1×10^6^ T cells were injected, and the cells were checked prior to transplant for a 1:1 ratio of CD4/CD8 for each strain, and for a 1:1 ratio of donor strains (**Fig. 6A**). As a control, we also transplanted WT CD45.2 (C57BL/6) and WT CD45.1 (B6-Ly5.1) cells at 1:1 ratio of WT(CD45.2): WT(CD45.1). Once again 1×10^6^ T cells were injected and cells were checked prior to transplant for a 1:1 ratio of CD4/CD8 for each strain, and for a 1:1 ratio of donor strains (**Fig. 6A**). At 7 days post transplantation, recipient mice were examined for the presence of donor T cells in the spleen, liver and small intestines.

**Figure 6.**
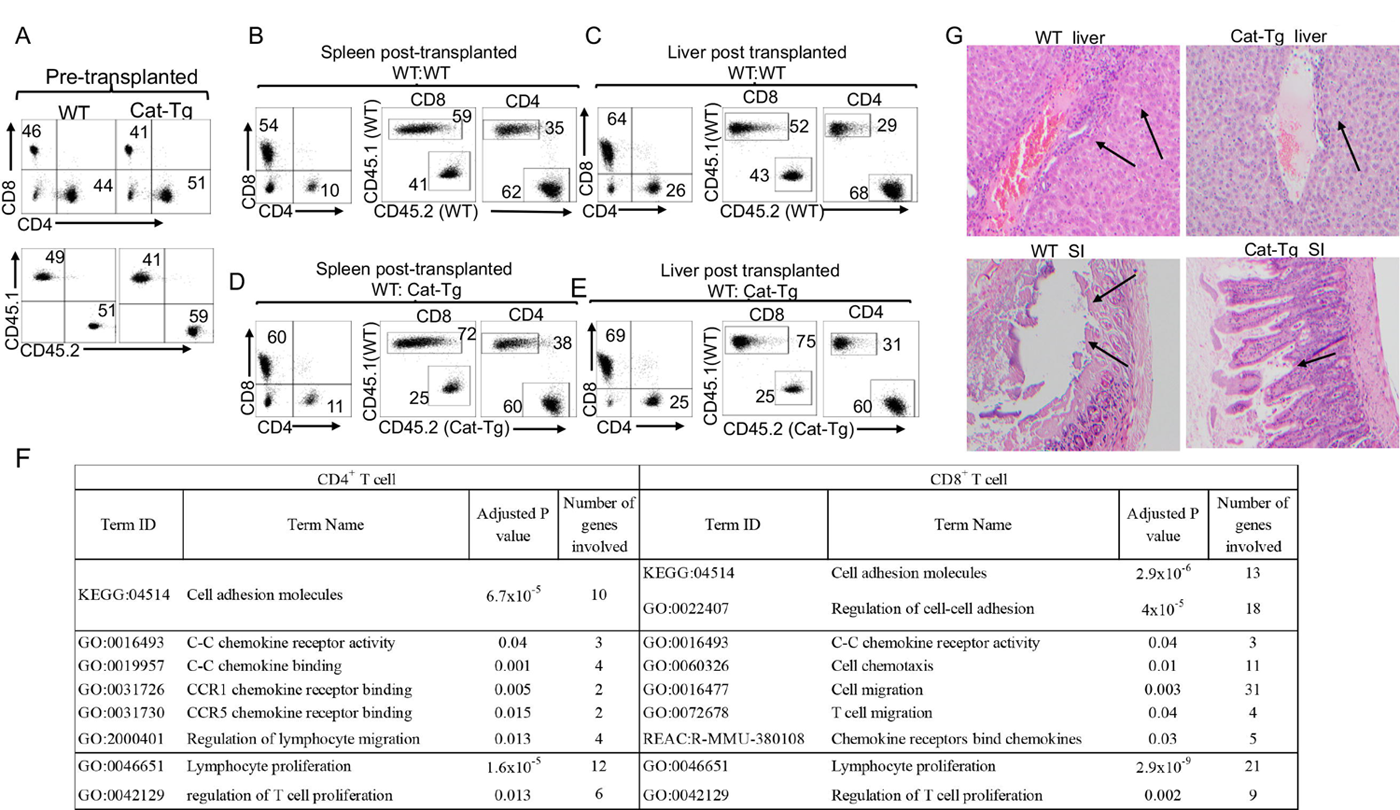
Wnt/β-catenin over expression specifically affects CD8^+^ T cell functions. **(A)** Irradiated BALB/c mice were allo-transplanted and injected with MACS-sorted WT and *Cat-Tg* CD4^+^ or CD8^+^ T cells mixed at a 1:1 ratio. Flow analysis of MACS sorted pre-transplanted T cells, showing an approximate 1:1 ratio of CD4:CD8 per strain, and an approximate 1:1 ratio of WT(CD45.1): *Cat-Tg* (CD45.2): cells (or WT(CD45.1): WT(CD45.2) as control). **(B)** At day 7 post-BMT, the spleen was examined for donor WT(CD45.1): WT(CD45.2) T cells. The percentage of WT(CD45.1): WT(CD45.2) CD4^+^ and CD8^+^ T cells is shown, as well as the percentage of each donor cell type in spleen. **(C)** At day 7 post-BMT, the liver was examined for donor T cells. The percentage of WT(CD45.1): WT(CD45.2) CD4^+^ and CD8^+^ T cells is shown, as well as the percentage of each donor cell type in liver. **(D)** At day 7 post-BMT, the spleen was examined for donor WT(CD45.1): *Cat-Tg* (CD45.2) T cells. The percentage of WT(CD45.1): *Cat-Tg* (CD45.2) CD4^+^ and CD8^+^ T cells is shown, as well as the percentage of each donor cell type in spleen. **(E)** At day 7 post-BMT, the liver was examined for donor WT(CD45.1): *Cat-Tg* (CD45.2) T cells. The percentage of WT(CD45.1): *Cat-Tg (*CD45.2) CD4^+^ and CD8^+^ T cells is shown, as well as the percentage of each donor cell type in liver. **(F)**. Pathways involved in chemokine receptor activity, chemotaxis, migration, and proliferation from functional pathway analysis of CD4^+^ T cells (Fig. 4D) and CD8^+^ T cells (Fig 5D). Table compares adjusted p values and number of genes involved in each pathway between CD4^+^ and CD8^+^ T cells. **(G)** At day 7 post-allo-HSCT, small intestines and liver were examined by H&E staining for tissue damage. Arrows show lymphocyte infiltration and tissue damage. Statistical analysis was performed using one-way ANOVA. Mann Whitney U test, P value presented with the figure. Symbol meaning for P values are: ns - P > 0.05; * - P ≤ 0.05; ** - P ≤ 0.01; *** - P ≤ 0.001; **** - P ≤ 0.0001. (For Fig.6A-E, n = 5 mice per group, repeated twice, one representative is shown. For Fig.6F-G, n=5 mice used per group)

Our data show that recipient mice transplanted with WT CD45.1 and WT CD45.2 had significantly reduced donor CD4^+^ cells versus donor CD8^+^ T cells in the spleen and liver. Next, we examined transplanted CD4^+^ and CD8^+^ T cells by CD45.1 and CD45.2 and we did not observe significant differences among CD8^+^ T cells in spleen. We observed that donor CD4^+^ T cells were more skewed toward WT C57BL/6 cell (CD45.2) (**Fig. 6B, Sup. Fig.4A**). Next, we examine donor T cell migration into liver in recipient mice transplanted with WT CD45.1 or WT CD45.2 cells. We observed a similar effect on donor T cells migration as we observed in recipient spleen (**Fig. 6C, Sup. Fig. 4B**). We also examined donor T cell migration into small intestines of recipient mice transplanted with WT CD45.1 or WT CD45.2 cells. We did not observe differences in CD4^+^ to CD8^+^ donor T cells ratios, and also, we also did not observe significant differences in CD45.1/CD45.2 expression in CD8^+^ T cells either from small intestine. We did observe donor CD4^+^ T cells were more skew toward WT C57 cells (CD45.2+) **(Sup. Fig4C**). When we examined spleens from recipient mice transplanted with CD45.1 WT and CD45.2 *Cat-Tg* cells, we observed that the ratio of CD4^+^ and CD8^+^ T cell in the spleen was similar to that seen in CD45.1 WT: WT CD45.2 transplanted mice. However, we observed a significant reduction in donor CD8^+^ T cells from CD45.2 *Cat-Tg* mice in recipient mouse spleen (**Fig. 6D**, **Sup. Fig.4D**). We also observed that there was no significant migration defect on donor CD4^+^ T cells; however, we observed a significant decrease in CD8^+^ T cell migration to recipient liver (one of the GVHD target organs) (**Fig. 6E, Sup. Fig.4E**). We also looked at the donor T cell migration into small intestines (another target organ of GVHD), and found significantly reduced CD4^+^ T cells compared to CD8^+^ T cells from donors after transplantation. There was a significant reduction in CD8^+^ donor T cells from *Cat-Tg* (CD45.2) mice in recipient spleen, and we observed that donor CD4^+^ T cells were more skewed towards *Cat-Tg* cells (CD45.2) **(Sup. Fig.4F**). To investigate the underlying mechanism behind these changes, we performed pathway analysis using RNA seq as an unbiased approach. Our data confirmed that both CD4^+^ and CD8^+^ T cells are affected by Wnt/β-catenin over-expression. We observed that KEGG pathways including cell adhesion molecules (10 genes) were affected in CD4^+^ T cells from *Cat-Tg* mice compared to WT mice. Our data show that in CD8^+^ T cells from *Cat-Tg* mice compared to WT mice, in addition to the KEGG Cell adhesion molecules pathway, we also observed 18 genes regulating cell to cell adhesion pathways (from GO: BP source) that were differentially regulated. We also observed that C-C chemokine receptor activity, cell chemotaxis, and cell migration (specifically T cell migration genes) were more significantly affected in CD8^+^T cells (*Cat-Tg* vs. WT mice) than in CD4^+^T cells. Furthermore, we also observed that genes involved in lymphocyte and T cell proliferation were more significantly affected in CD8^+^T cells (30 genes) than in CD4^+^ T cells (18 genes) (**Fig. 6F**).

Using histological staining for H&E, we also observed leukocyte infiltration into GVHD target organs like liver and small intestine (SI)(33) in WT T cell recipients, but not as much in *Cat-Tg* T cell recipients (**Fig. 6G**) and (**Supp. Fig.5**). These data suggest that CD8^+^ T cells from *Cat-Tg* mice have significantly been affected by β-catenin over-expression, and caused less tissue damage to GVHD target tissues.

## Discussion

In this report, we demonstrate that over-expression of Wnt/β cells for the treatment of hematological malignancies. Both CD4^+^ and CD8^+^ T cells from mice over-expressing β−catenin (*Cat-Tg* mice) showed significantly reduced GVHD pathogenesis, while maintaining GVL in models of allo-HSCT. CD8^+^ and CD4^+^ T cells from *Cat-Tg* mice expressed higher levels of CD44 and PD-1 markers. CD8^+^ T cells from *Cat-Tg* mice also expressed higher Eomes and CD122. We did not observe any differences in TCF-1, T-bet or CTLA-4 expression. CD8^+^ T cells from *Cat-Tg* mice showed frequencies of central memory and transitioning cells, but a reduced naïve cell population.

Several lines of evidence suggested that CD44^hi^ and CD122^hi^ T cells do not induce GVHD(36–38). Our data showed that a high proportion of CD8^+^ T cells from *Cat-Tg* mice are CD44^hi^ and CD122^hi^ and express higher levels of Eomes (IMP phenotype) (29). Previously, it has been suggested that the IMP phenotype might be due to higher expression of IL-4 in the in the thymus of *Cat-Tg* mice, which can result in the IMP phenotype (39). However, published data have indicated that the IMP phenotype is not dependent on IL-4 expression specifically in *Cat-Tg* mice (8, 40). These findings would indicate that higher Eomes expression and the IMP phenotype in *Cat-Tg* mice due to β−catenin over-expression allows these cells to have anti-tumor activity in a T cell-intrinsic manner.

Several lines of evidence also suggest that β-catenin plays a central role in T cell development (12, 41, 42). Experiments using either loss of β-catenin or enforced expression of stabilized β-catenin have further identified a role for β-catenin at multiple stages of T cell development (42, 43). Adoptive transfer of Wnt-treated CD8^+^ T cells was shown to enhance anti-tumor activity *in vivo* (44). Our data provide evidence that CD8^+^ T cells from *Cat-Tg* mice express higher levels of granzyme B and perforin, and higher expression of Eomes, and these cells also exhibited enhanced cytotoxicity. Constitutive activation of the TCF-1/β pathway *in vivo* has been shown to favor generation of memory CD8^+^ T cells (45).

To examine how T cells from *Cat-Tg* mice maintain GVL function without GVHD damage, we examined proinflammatory cytokine expression. Our data show that donor T cells from *Cat-Tg* mice express significantly less proinflammatory cytokines both on a serum level and on a cellular level in our allo-HSCT model. Over-expression of β−catenin did not alter the signaling molecules ERK, PLCγ1, and IRF-4. Wnt/β catenin signaling, a highly conserved pathway through evolution, regulates key cellular functions including proliferation, differentiation, migration, genetic stability, apoptosis, and stem cell renewal (43, 45). Donor T cells were examined for proliferation using an EdU incorporation assay. Both donor CD4^+^ and CD8^+^ T cells from *Cat-Tg* mice showed a trend toward reduced proliferation compared to donor T cells from WT mice. To investigate how the Wnt/β expression, we utilized an unbiased RNA sequencing approach. Transcriptome analysis by RNA sequencing revealed that there were 150 differentially expressed genes in CD4^+^ T cells **(Sup. Table 1**), and over 250 genes affected by over-expression of β−catenin in CD8^+^ T cells **(Sup. Table 3**). Pathway analysis revealed that the differentially expressed genes in CD4^+^ T cells are involved in regulation of immune system processes, T cell and B cell activation, T cell proliferation, adaptive immune responses, immune system development, inflammatory responses, cytokine production, signaling, cell adhesion, and chemokine receptors. Genes that were differentially expressed in CD8^+^ T cells were involved in similar pathways, along with hematopoietic cell lineage, GVHD, allograft rejection, Th1, Th2, Th17 differentiation, and other pathways. Therefore, β−catenin plays a critical role in regulating gene expression programs of mature T cells during alloactivation. Taken together, these data suggest that Wnt/β represent a potential target for the separation of GVHD and GVL responses after allo-HSCT. Next, we sought to investigate how these transcriptome changes affected donor T cell function in the development of GVHD pathophysiology. Donor T cell proliferation and migration to GVHD target organs are considered hallmarks of GVHD (4). Therefore, we examined whether T cells from *Cat-Tg* are defective in migration to GVHD target organs. We observed that only CD8^+^ T cells from *Cat-Tg* mice are likely to be defective. This effect could also be due to the observed decreased in proliferation or decreased cell survival, but alteration on gene programs related to migration suggested that Cat-Tg CD8+T cells are at least partly migration defective. When transplanting a 1:1 ratio of WT: *Cat-Tg* T cells, we did not observe any differences in migration of CD4^+^ T cells from *Cat-Tg* mice compared to WT mice in spleen and liver. We observed a significant reduction in CD8^+^ T cells from *Cat-Tg* mice in spleen. Similarly, we observed a significant reduction in CD8^+^T cells from *Cat-Tg* mice in liver and small intestines as well. We observed that donor T cells from *Cat-Tg* mice trend towards decreased proliferation. We also examined genes involved in total lymphocyte proliferation and specifically genes involved in T cell proliferation, which were affected in both CD4^+^ and CD8^+^ T cells, but significantly more genes were affected in donor CD8^+^T cells compared to CD4^+^ T cells from *Cat-Tg* mice.

Activation-induced cell death (AICD) of lymphocytes is an apoptotic pathway that might be involved in the control of CD8^+^ T cell homeostasis in *Cat-Tg mice* (46). Therefore, we performed pathway analysis, and our data confirm that many pathways were significantly affected in both CD4^+^ and CD8^+^ T cells from *Cat-Tg* mice. Our data confirmed that cell adhesion molecules were more significantly affected in CD8^+^ T cells than in CD4^+^T cell from *Cat-Tg* mice. We also observed a significant reduction in cell adhesion molecule expression (13 genes) cell adhesion molecule expression (18 genes) in CD8. Published data have shown that irradiation causes the upregulation of cell adhesion molecules and provides early costimulatory signals to incoming donor T cells in the intestine, followed by a cascade of proinflammatory signals in other organs once the alloresponse is established (47). Our data provide evidence that both CD4 and CD8^+^T cells from *Cat-Tg* mice show significant reductions in adhesion molecules, but CD8^+^ T cells from *Cat-Tg* mice were more significantly affected.

We also examined pathways involved in lymphocyte migration by pathway analysis. We observed that genes involved in chemokine receptor activity, C-C chemokine binding, CCR1 chemokine receptor binding, CCR5 chemokine receptor binding, and several genes involved in the regulation of lymphocyte migration were affected in CD4^+^ T cells. More genes in similar pathways like C-C chemokine receptor activity, Cell chemotaxis, Chemokine receptors binding chemokines, and especially Cell migration and T cell migration were found to be significantly altered in CD8^+^ T cells from *Cat-Tg* mice compared to WT mice. Inflammatory chemokines are expressed in inflamed tissues by both hematopoetic and non-hematopoetic cells upon stimulation by pro-inflammatory cytokines, including TNFα and IFN-γ (48). Inflammatory chemokines of the CC, C, or CXC3C families are also increasingly expressed after allogeneic transplantation (48). Cellular sources of chemokines may differ between specific target organs, and contribute considerably to the severity of GVHD. Our data show that inflammatory chemokines are significantly affected in T cells from *Cat-Tg* mice, which contributed to less severe GVHD development. Our model demonstrated that T cells from *Cat-Tg* mice significantly reduce the development of acute GVHD.

Recently, several lines of evidence have suggested that modulating intracellular signaling pathways that regulate T cell responses and survival can be used to inhibit T cell alloresponses, T cell survival, and thus, GVHD (49). This has included targeting the transcription factor nuclear factor kappa B (NFκB). As NFκB has long been known to play a critical role in T cell biology, particularly with respect to cytokine responses, it has always been an attractive target (50).

However, targeting of NFκB has been hampered by a lack of reagents that have favorable pharmacokinetics, resulting in systemic NFκB inhibition without toxicity(51, 52). Our data show that T cells (both CD4 and CD8^+^) from *Cat-Tg* mice have significant changes to the NFκB pathways.

NFκB is a transcription factor that controls the expression of a number of genes important for mediating immune and inflammatory responses. Several lines of evidence have recently suggested that inhibiting NFκB signaling ameliorates GVHD in both mice and humans (53). Bortezomib and PS-1145 are small molecule inhibitors that have been used to treat acute GVHD (54). In our RNAseq results, we also observed a number of B cell activation and regulation pathways, as well as immunoglobulin regulation pathways, which were significantly affected by β-catenin over-expression in both CD4 and CD8+ T cells. Even though we didn’t investigate β-catenin impacts B cells in GVHD pathogenesis at a functional level, it β has been previously shown that B cells play an important role in pathogenesis of GVHD(55), so this could be another reason why over-expression of β-catenin signaling will have considerable clinical β implications for the improvement of immunotherapies based on *ex vivo* manipulation of T lymphocytes for adoptive transplantation. Several mouse models have shown that blocking Gsk-3β using small molecule inhibitors resulted in the generation of stem-like memory CD8^+^ T cells, which have the potential to be highly effective in immunotherapy (56). Using pharmacological approaches, human CD8^+^ T cells with stem-like properties can be generated using antagonists of Wnt signaling, and can be genetically engineered to have tumor-specific properties (57). For the first time we showed that Wnt/β-catenin is not only critical for T cell development, but also plays a significant role in T cells-mediated GVHD after allogeneic transplantation. Our functional and genetic data demonstrated that the Wnt/β-catenin pathways play a central role in uncoupling GVHD from GVL functions.

## Supporting information

FDP

## Acknowledgements

We thank all members of the Karimi laboratory for helpful discussions. This research was funded in part by a grant from the National Blood Foundation Scholar Award to (MK), the National Institutes of Health (NIH LRP #L6 MD0010106 and K22 (AI130182) to MK), and an Upstate Medical University Cancer grant (1146249-1-75632) to MK. JMS and AV are supported by the Intramural Research Program of the National Institute of Aging.

## Author contributions

MM, RH, SM, AM, RD, and MK performed experiments; JMS provided valuable reagents; and helped edit the manuscript. AV, provided reagents. AW helped with RNA sequencing analysis. MM, RH, and MK designed experiments, analyzed the data, and wrote the manuscript.

## Supplementary Figure Legends

Supplementary Figure 1. **T cells from Cat-Tg mice exhibit enhanced T cell IMP phenotypes, related to** Figure 2**. (A-B)** Quantitative analysis of T cells from WT and *Cat-Tg* mice that were examined for effector memory, central memory, transitioning/activating, and naïve population frequencies. CD4 **(A)** and CD8 **(B)** T cells were examined for these populations by flow cytometry. Statistical analysis was performed using two-way ANOVA, one-way ANOVA confirmed by Student’s *t-*test, p-values are presented. Symbol meanings for P-values are: ns - P > 0.05; * - P ≤ 0.05; ** - P ≤ 0.01; *** - P ≤ 0.001; **** - P ≤ 0.0001(n = 3 mice per group)

Supplementary Figure 2. **GSEA enrichment analysis for CD4^+^ T cells, related to** Figure 4**. (A)** A bubble chart showing up- or downregulated pathways based on enrichment scores in GSEA in pre-transplanted WT or *Cat-Tg* CD4^+^ T cell samples. **(B)** A bubble chart showing up- or down regulated pathways based on enrichment scores in GSEA in day 7 post-transplant WT or *Cat-Tg* CD4^+^ T cell samples.

Supplementary Figure 3. **GSEA enrichment analysis for CD8^+^ T cells, related to** Figure 5**. (A)** A bubble chart showing up- or downregulated pathways based on enrichment scores in GSEA in pre-transplanted WT or *Cat-Tg* CD8^+^ T cell samples. **(B)** A bubble chart showing up- or down regulated pathways based on enrichment scores in GSEA in day 7 post-transplant WT or *Cat-Tg* CD8^+^ T cell.

Supplementary Figure 4. **Wnt/**β **catenin over-expression specifically affects CD8 T cell functions, related to** Figure 6**. (A-C)** Quantitative analysis using flow cytometry of donor CD8^+^ and CD4^+^ T cells isolated from recipient’s spleen, liver, and small intestine (SM) seven days post transplantation. Donor CD4^+^ and CD8^+^ T cells were further analyzed by CD45.1 (WT-B6-Ly5.1) and CD45.2 (WT control-C57BL/6) to identify donor strains. **(D-F)** Quantitative analysis using flow cytometry of donor CD8^+^ and CD4^+^ T cells isolated from recipient spleen, liver, and small intestine (SM) seven days post transplantation. Donor CD4^+^ and CD8^+^ T cells were further analyzed by CD45.1 (WT) and CD45.2 (Cat-Tg) to identify donor strains. For statistical analysis we used one-way ANOVA and student’s *t* test, *p* values are presented. Symbol meaning for P values are: ns - P > 0.05; * - P ≤ 0.05; ** - P ≤ 0.01; *** - P ≤ 0.001; **** - P ≤ 0.0001(n = 5 mice per group, repeated twice, one representative is shown).

Supplementary Figure 5. **Quantified scores for GVHD target organ damage, related to** Figure 6 Tissues obtained from recipient mice were graded for GVHD. Quantified GVHD scores for different groups are shown, with median indicated.. Statistical analysis was performed using Mann Whitney U test.

