## Supplementary material for "Wnt/β-catenin regulates alloreactive T cells for the treatment of hematological malignancies": FDP

#### Supplementary Figure 1

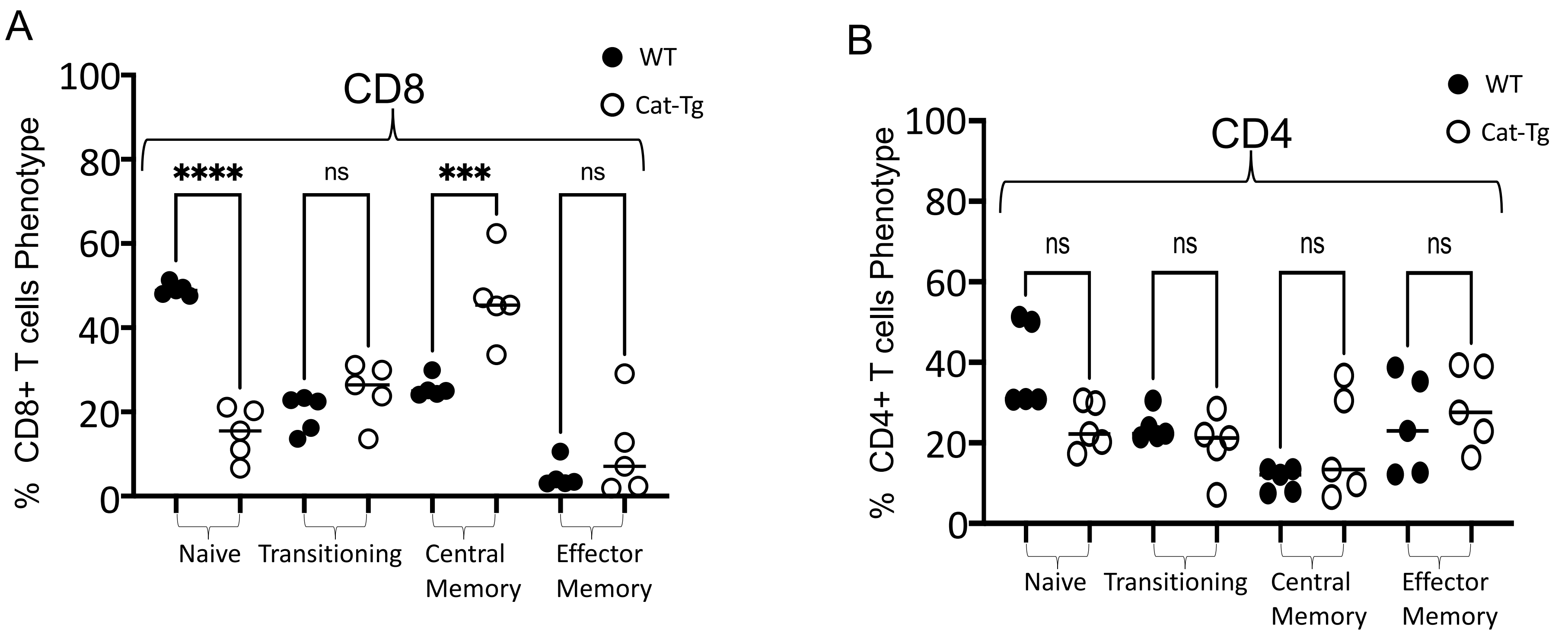

### Supplementary Figure 2

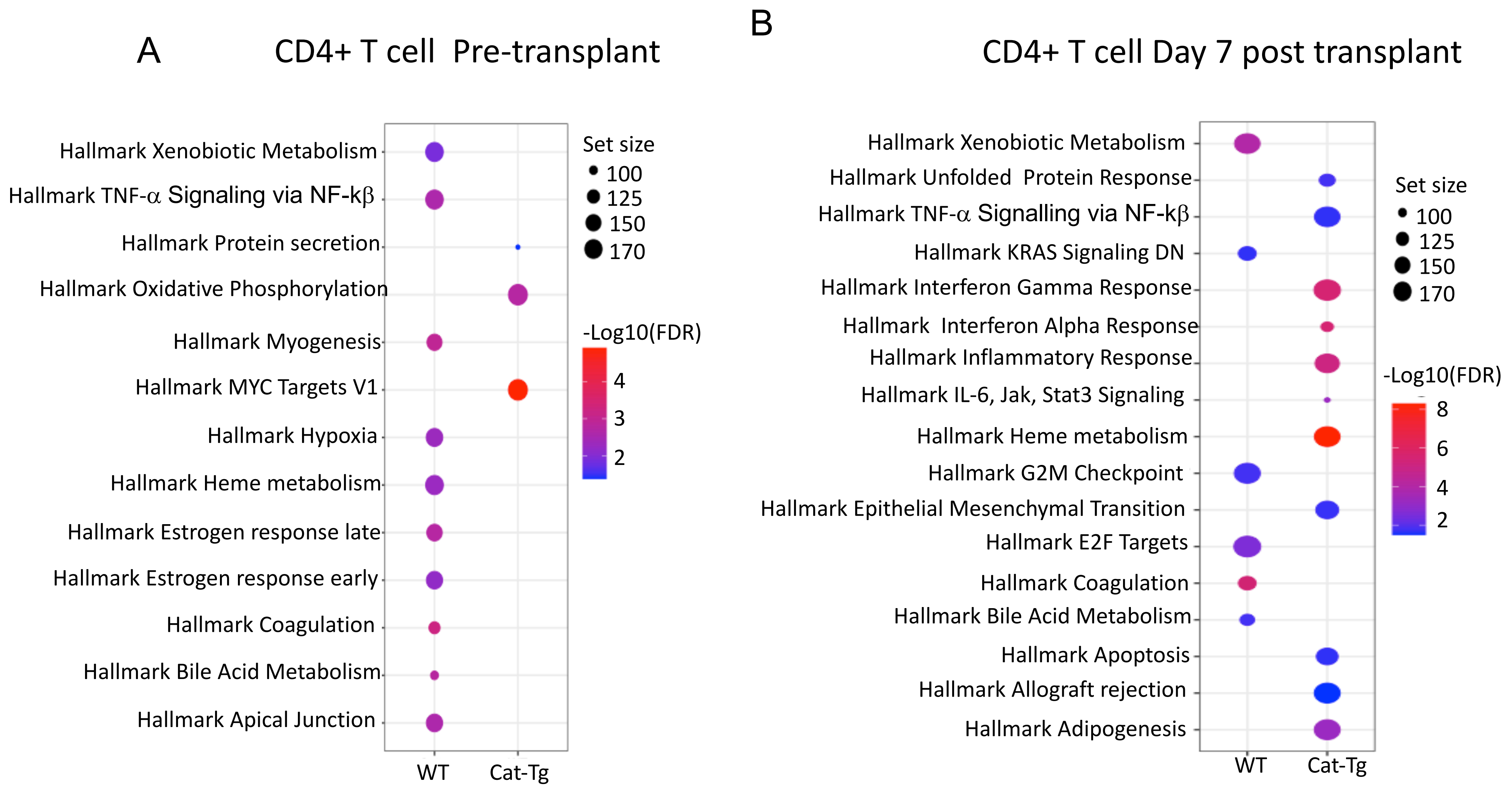

Supplementary Figure 3

A

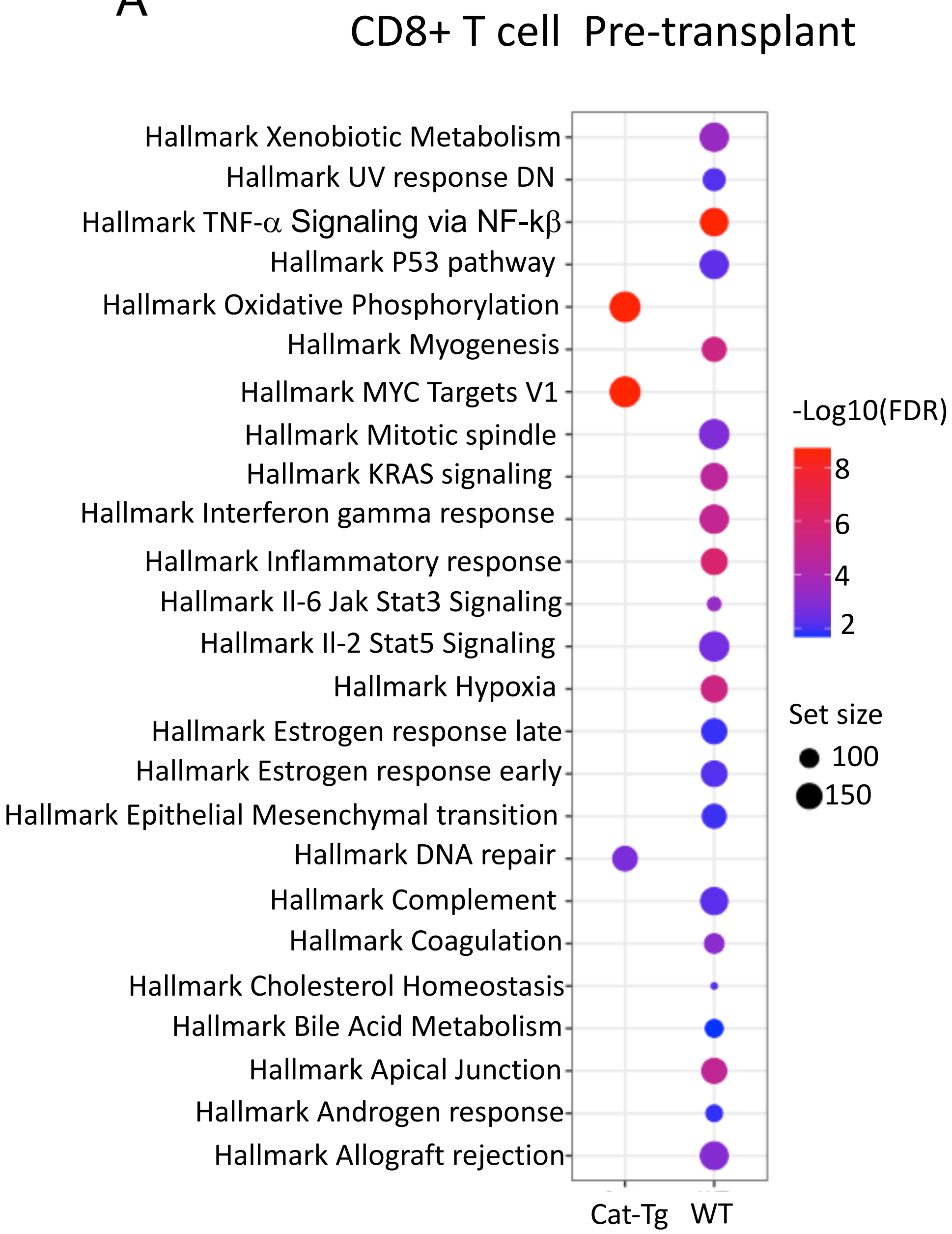


B

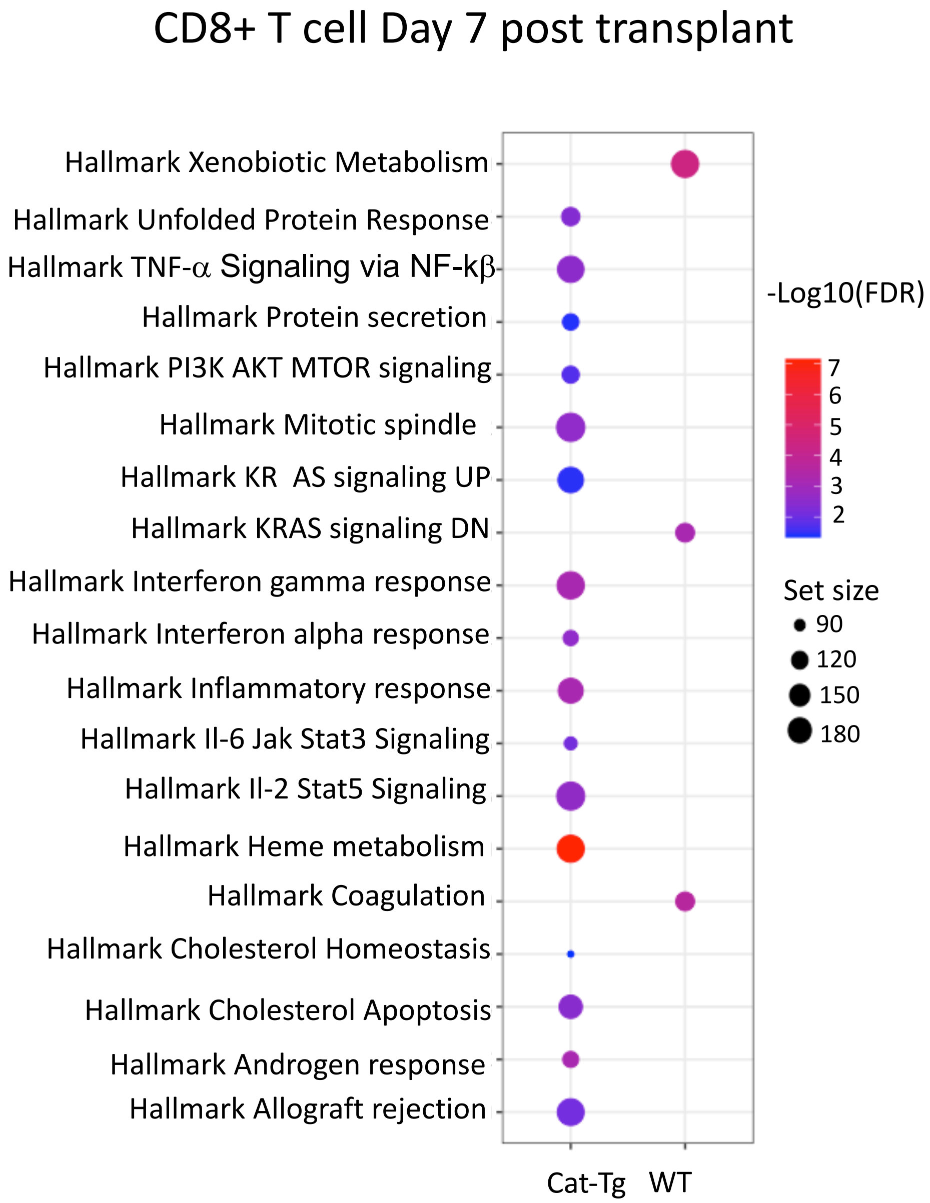

### Supplementary Figure 4

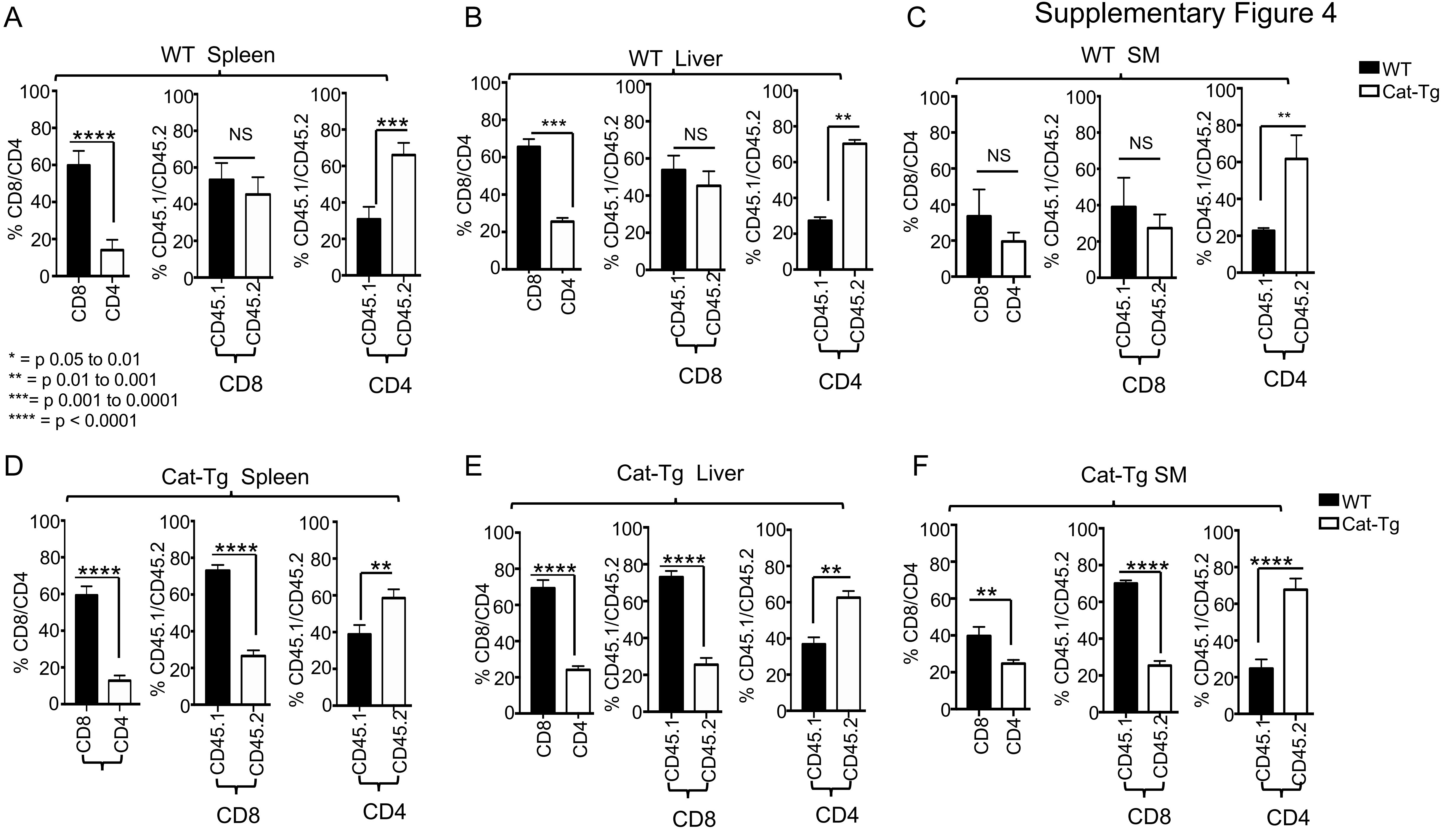

Supplementary Figure 5

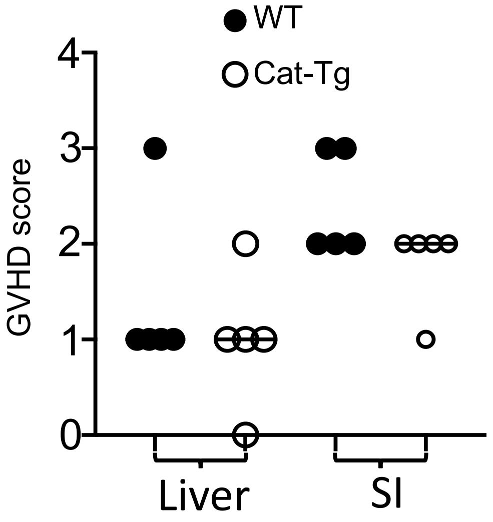
